# Phage intervention improves colitis and response to corticosteroids by attenuating virulence of Crohn’s disease-associated bacteria

**DOI:** 10.1101/2025.06.05.658057

**Authors:** Kyle Jackson, Heather Galipeau, Amber Hann, Marco Constante, Megan Zangara, Michael Bording-Jorgensen, Alexandra Fuentes, Heidi Ho, Jessica Wang, Chiko Shimbori, Paul Moayyedi, Michael Surette, Premysl Bercik, Brian Coombes, Zeinab Hosseinidoust, Elena F. Verdu

## Abstract

Adherent-invasive *Escherichia coli* (AIEC) exhibit proinflammatory properties and have been implicated in the pathogenesis of Crohn’s disease (CD), a form of inflammatory bowel disease (IBD). Antibiotic use in CD lacks specificity and may worsen microbiome disruption, prompting interest in bacteriophages (phages) for targeted microbiome editing. Here, we identified HER259, a phage active against the clinical AIEC strain NRG857c. Using gnotobiotic models of AIEC-driven colitis, we show that HER259 attenuates AIEC virulence, including suppression of the FimH adhesin through inversion of the *fimS* promoter to its ‘off’ orientation. Withdrawal of HER259 treatment leads to reversion of the *fimS* promoter and reactivated colitis in mice. HER259 phage also enhances the therapeutic effect of sub-therapeutic budesonide, independent of microbial drug metabolism. These findings support targeted phage therapy as an adjunct treatment approach in IBD, demonstrating modulation of bacterial virulence and improved response to conventional treatments which may reduce drug-related side effects.

**One Sentence Summary:** Bacteriophage HER259 improves colitis severity mediated by Crohn’s disease *Escherichia coli* NRG857c, and increases efficacy of budesonide.

## Introduction

Inflammatory bowel diseases (IBD) are chronic inflammatory conditions of the gastrointestinal (GI) tract that include Crohn’s disease (CD) and ulcerative colitis. Their pathophysiology is multifactorial, but is linked to dysregulated environment-host interactions.(*1–3*) Indeed, recent studies have demonstrated that changes in the composition and function of the gut microbiota precede CD onset by up to five years.(*4*) Despite this, current medications focus on suppressing symptoms and inflammation, and can fail, in part, because they do not address underlying microbial drivers.(*5, 6*)

Antibiotics are often prescribed during disease flares but are not ideal, given their lack of specificity and the adverse events associated with their frequent use.(*7*) These include risks of developing a *Clostridium difficile* infection(*8*) and the amplification of microbial perturbations as associated with disease onset.(*9*) For these reasons, attention has turned to bacteriophages (phages), which are ubiquitous self-replicating bacterial viruses that display high specificity and selectivity against their targeted bacteria. Indeed, commercial phage “cocktails” are presently being evaluated in clinical trials (NCT04737876, NCT03808103) against proinflammatory bacterial taxa associated with IBD. Specifically, these trials seek to target adherent-invasive *Escherichia coli* (AIEC), a group of bacteria that are enriched in patients with active CD compared with those in remission, or healthy individuals.(*10*) AIEC are distinct from other IBD-associated taxa by their ability to adhere, invade and survive in intestinal epithelial cells.(*11*) Adherence and invasion are mediated, in part, by the type 1 fimbriae D-mannose specific adhesin (FimH) governed by the *fimS* ON/OFF switch.(*12*) In the “ON” orientation, the 314-bp fimS element aligns the promoter to drive transcription of the *fim* operon, including *fimH*. This ability is considered a key virulence mechanism that can exacerbate inflammation in CD, whose orientation can be influenced by environmental signalling.(*13, 14*) Thus, approaches to target AIEC in a specific manner using bacteriophages, have garnered interest.(*15, 16*) However, primary outcomes for evaluating successful phage intervention in general emphasize the reduction in global bacterial load and eradication, which neglects nuances in microbe-host interactions in IBD, where elimination of a taxon could worsen an already perturbed microbial community.

Thus, our aim was to elucidate mechanisms that underpin phage therapy efficacy in colitis. Using well-characterized CD-associated *E. coli* strains, we evaluated their ability to worsen gut inflammation in chemical and genetic models of colitis in gnotobiotic conditions. We identified phage HER259 that specifically targets the most colitogenic strain. Targeted phage intervention modulated bacterial virulence without eradication, leading to significant clinical improvements and increased responses to corticosteroid budesonide. Based on our findings, targeted suppression of bacterial virulence by phage emerges as a promising new strategy for colitis control.

## Results

### Distinct E. coli with pro-inflammatory potential drive colitis severity in gnotobiotic mice

We developed gnotobiotic mouse models where germ-free (GF) C57BL/6 mice were *de novo* co-colonized with an Altered Schadler flora-like (ASF) community lacking Proteobacteria (**Fig S1**) and one of three AIEC strains, *E. coli* C0004 (urinary isolate), *E. coli* LF82 (CD isolate), and *E. coli* NRG857c (CD isolate).(*17, 18*) After 3 weeks of microbiota stabilization, we assessed their individual colitogenic capacity in the acute injury model of low-grade dextran sodium sulfate (DSS) (2% w/v) (**Fig 1A**). Control mice, colonized in the same manner, received water alone (**Fig S2A**). Prior to colitis induction, 16S rRNA gene sequencing confirmed that *E. coli* was only present in mice *de novo* colonized with the AIEC strains (**Fig S1B**). Mice colonized with *E. coli* had more severe colitis than those with ASF alone, but the severity varied by *E. coli* strain (**Fig 1B, 1C**). ASF+NRG857c mice had more severe clinical indicators of colitis, including loose stool consistency and fecal occult blood two days earlier than mice with other *E. coli* isolates (**Fig 1B**). While disease activity index (DAI) at endpoint was similar between mice colonized with ASF+NRG857c or ASF+LF82, the area under the curve (AUC) for DAI throughout the monitoring period was higher in ASF+NRG857c compared with ASF+LF82-colonized mice (**Fig 1C**, p<0.01). Histological scores were also higher in ASF+LF82 or ASF+NRG857c than ASF+C0004, the latter a urinary isolate (**Fig 1D, 1E**, p<0.01). Differences in colitogenic capacity between strains were independent of cecal bacterial load at endpoint (**Fig 1F**). Mice that did not receive DSS had no signs of colitis, irrespective of colonization with *E. coli* (**Fig S2B, S2C**). Moreover, there were no significant changes in beta diversity before and after DSS treatment across all groups (**Fig S1E-F**). Based on the collective results, we selected the most colitogenic AIEC strain, NRG857c, for subsequent experiments using the spontaneous chronic colitis model in interleukin-10 deficient (*IL-10^-/-^*) mice.

**Fig. 1.**
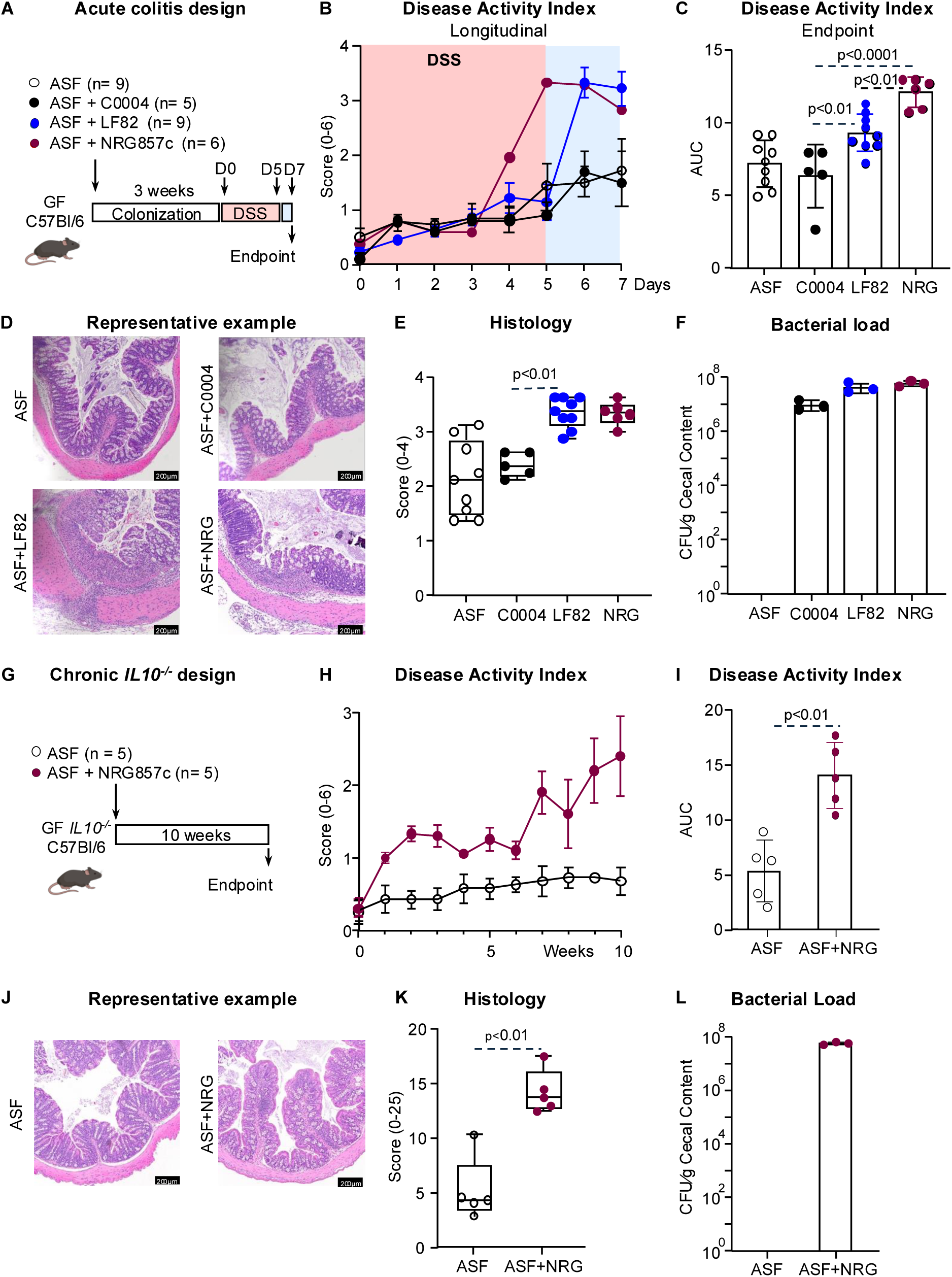
CD-associated AIEC exacerbate colitis in gnotobiotic mouse models. (**A**) Germ-free C57Bl/6 mice were co-colonized with ASF and 10^8^ CFU of *E. coli* NRG857c, for 3 weeks. Mice were treated with low grade DSS (2% w/v) for five days followed by two days of water. (**B**) Disease activity index (stool consistency + occult blood) analysed longitudinally. (**C**) Area under the curve for disease activity index at endpoint. (**D**) Representative histology examples of cross-sectional proximal colon (20X). (**E**) Histology scores (Cooper score method). (**F**) Cecal bacterial load at endpoint. (**G**) Germ-free C57Bl/6 IL-10^-/-^ mice were co-colonized with ASF and 10^8^ CFU of *E. coli* NRG857c and monitored weekly for 10 weeks. (**H**) Disease activity index (stool consistency + occult blood) analysed longitudinally. (**I**) Area under the curve for disease activity index at endpoint. (**J**) Representative histology examples of cross-sectional proximal colon (20X). (**K**) Histology scores, using a modified grading system. (**L**) Cecal bacterial load at endpoint. Statistical significance determined by Kruskal-Wallis test with Dunn’s post-hoc test or ANOVA with Tukey post-hoc test.

GF *IL-10^-/-^* mice were colonized with ASF and NRG857c (**Fig 1G**), and additional mice colonized with ASF alone served as controls. At 10-weeks post colonization, mice with NRG857c had higher clinical (**Fig 1H, 1I**) and histological scores compared with control mice at endpoint (**Fig 1J, 1K**, p<0.01). Long-term colonization of *IL-10^-/-^* mice with ASF+NRG857c was confirmed by bacterial load at sacrifice (**Fig 1I**). Altogether, these data showed that NRG857c exacerbates colitis in two distinct gnotobiotic animal models.

### Lytic phage controls bacterial load and decreases biofilms in vitro

We screened for phages capable of infecting NRG857c, LF82, and/or C0004 using the phages available at the Felix d’Herelle Reference Center for Bacterial Viruses at the Université Laval. Initial screens identified four T4-like lytic bacteriophages originally isolated for strain typing against *E. coli* O157:H7, namely HER256, HER258, HER259, and HER262 (**Fig 2A, Fig S3A-D**).(*19*) To assess the ability of these phages to produce a clearing on a bacterial lawn, we first performed spot tests (**Fig 2A**). All four phages had the capacity to target the bacterial strains of interest, as indicated by clearing on the bacterial lawn (**Fig 2A**). However, considering data presented in Fig 1, we focused on the two *E. coli* strains with the greatest colitogenic capacity: NRG857c and LF82. To further narrow down our phage selection, we compared the viral titer of each phage against the targeted bacterium to assess relative infectivity. The results revealed that HER259 and HER262 exhibited significantly higher viral titers against NRG857c and LF82 compared with HER256 and HER258, indicating an enhanced lytic activity towards these bacterial strains (**Fig 2B)**. In liquid culture kill curve assays of NRG857c or LF82, phages HER259 and HER262 showed marked differences in performance (**Fig 2C**). We also tested *in vitro* various multiplicities of infection (MOI) to determine the optimal dose for translation to our animal models. Phage HER259 exhibited the most effective bacterial suppression of NRG857c growth at an MOI of 10 (equivalent to 10⁹ PFU/dose) (**Fig 2C, top-left**). We also explored whether a combined regimen of HER259 and HER262 could enhance the targeted killing capacity against LF82 or NRG857c, motivated by the preference to use phage cocktails in clinical trials investigating phage therapy against AIEC. (*20*) We thus formulated a cocktail of HER259 and HER262 at a final concentration of MOI 10 and conducted *in vitro* kill curve assays against both strains (**Fig S4**). We observed that the cocktail interventions did not improve the bacterial load reduction against LF82 (**Fig S4A**) or NRG857c (**Fig S4B**) *in vitro*. Therefore, we concluded that the most effective phage-bacterial pairing was phage HER259 against *E. coli* NRG857c.

**Fig. 2.**
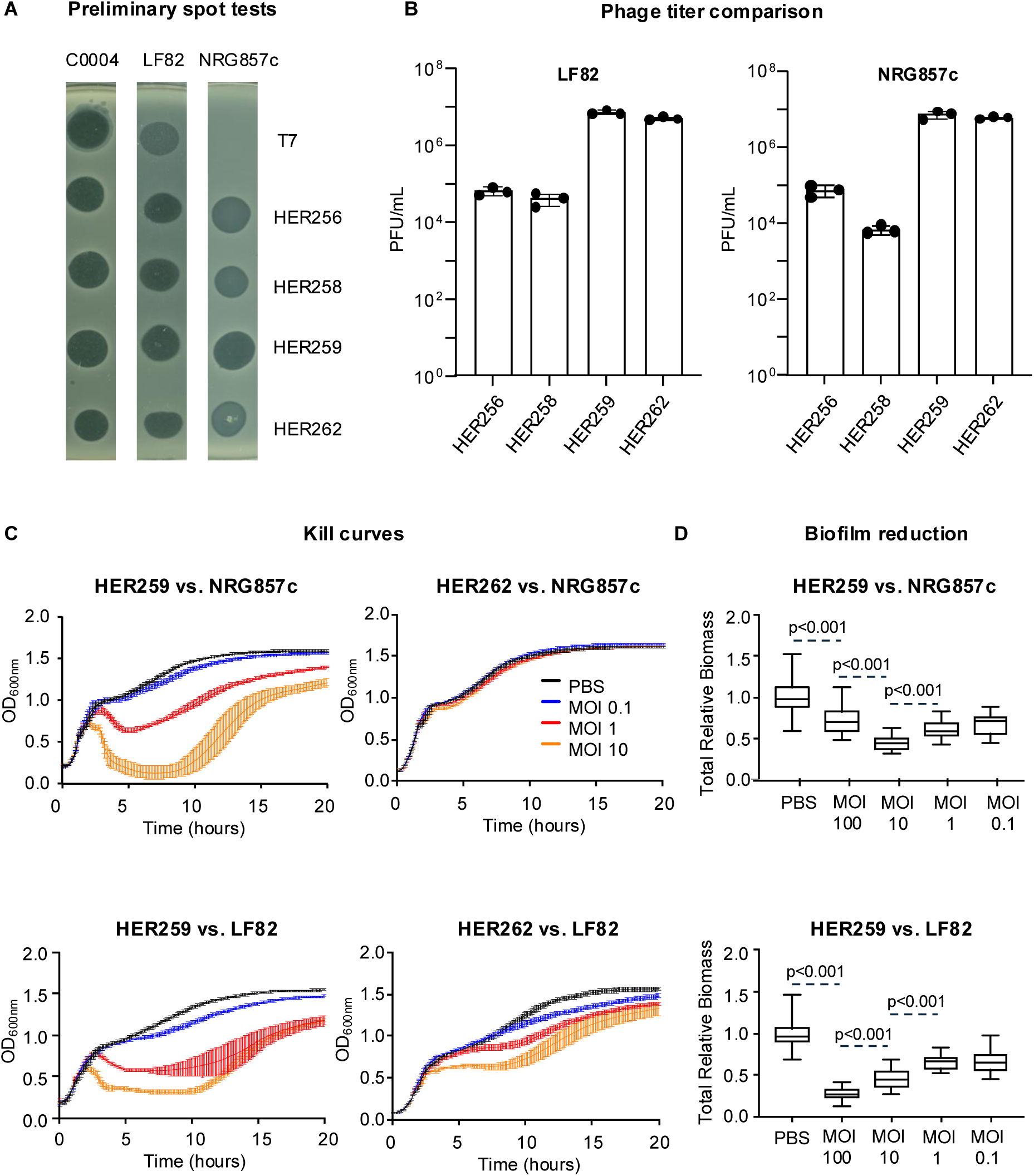
HER259 targets CD-associated AIEC. **(A**) Spot test of selected phage against C0004, LF82, and NRG857c. (**B**) Phage titer comparison of HER257, HER258, HER259, and HER262 against LF82 and NRG857c. (**C**) Phage kill curves of HER259 (left) and HER262 (right) against NRG857c (top) and LF82 (bottom). (**D**) Phage biofilm challenge of HER259 against NRG857c (top) and LF82 (bottom). Statistical significance determined by Kruskal-Wallis test with Dunn’s post-hoc test or ANOVA with Tukey post-hoc test.

To further enhance the infectivity of HER259 against NRG857c, we used an *in vitro* selection process by serially passaging the phage over multiple generations (**Fig S5A**). (*21*) After 10 cycles of passaging, we observed a 2-log increase in the viral titer of HER259 against NRG857c (**Fig S5B)**. This was complemented by an enhanced ability to remove biomass when challenged against NRG857c biofilms grown overnight in 96-well plates (**Fig S5C**).(*22*) Collectively, these results confirmed successful enhancement of infectivity against NRG857c. The passaged HER259 was used for further *in vitro* validation and subsequent *in vivo* testing.

To validate the optimal dosing of HER259 against NRG857c, we tested the passaged phage against NRG857c biofilms grown overnight in 96-well plates at MOIs of 100, 10, 1, and 0.1. The results indicated that an MOI of 10 was the most effective at reducing the total relative biomass (**Fig 2D**, top). We observed that biofilms formed by NRG857c did not increase in relative biomass between 24 and 48 hours of growth, indicating the reduction observed following HER259 challenge was due to the phage’s ability to decrease biomass rather than strictly inhibiting further growth (**Fig S6**). Based on the combined findings, we selected HER259 and NRG857c as a model phage-host pair, and a MOI of 10 as the optimal dosing concentration, to test in our gnotobiotic models.

### HER259 attenuates acute colitis in NRG857c colonized mice

We orally treated ASF+NRG857c mice for two weeks prior to DSS with either vehicle (control), or HER259 (**Fig 3A**). To assess the effect of different dosing frequencies, mice received phage either daily or 3 times per week (3xweek). 16S rRNA gene sequencing was investigated in feces at day 0 (prior to phage intervention) and at day 14 (post phage intervention but before colitis induction) (**Fig S7A**). Non-significant reductions in the relative abundance of NRG857c, as well as beta diversity shifts, were observed in all groups on day 14 (**Fig S7B-F**), suggesting that targeted HER259 challenge did not substantially alter the background microbiota. Low-grade DSS was then administered. Irrespective of the dosing schedule, mice treated with phage had lower DAI scores as compared with controls (**Fig 3B, 3C** p<0.0001), although 3xweek phage performed better than daily phage (**Fig 3C**, p<0.03). This suggested more frequent phage administration did not add therapeutic benefit. Moreover, mice treated with vehicle had significantly higher histological scores compared with those treated with phage daily (**Fig 3D, 3E** p<0.001) or phage 3xweek (**Fig 3E**, p<0.01).

**Fig. 3.**
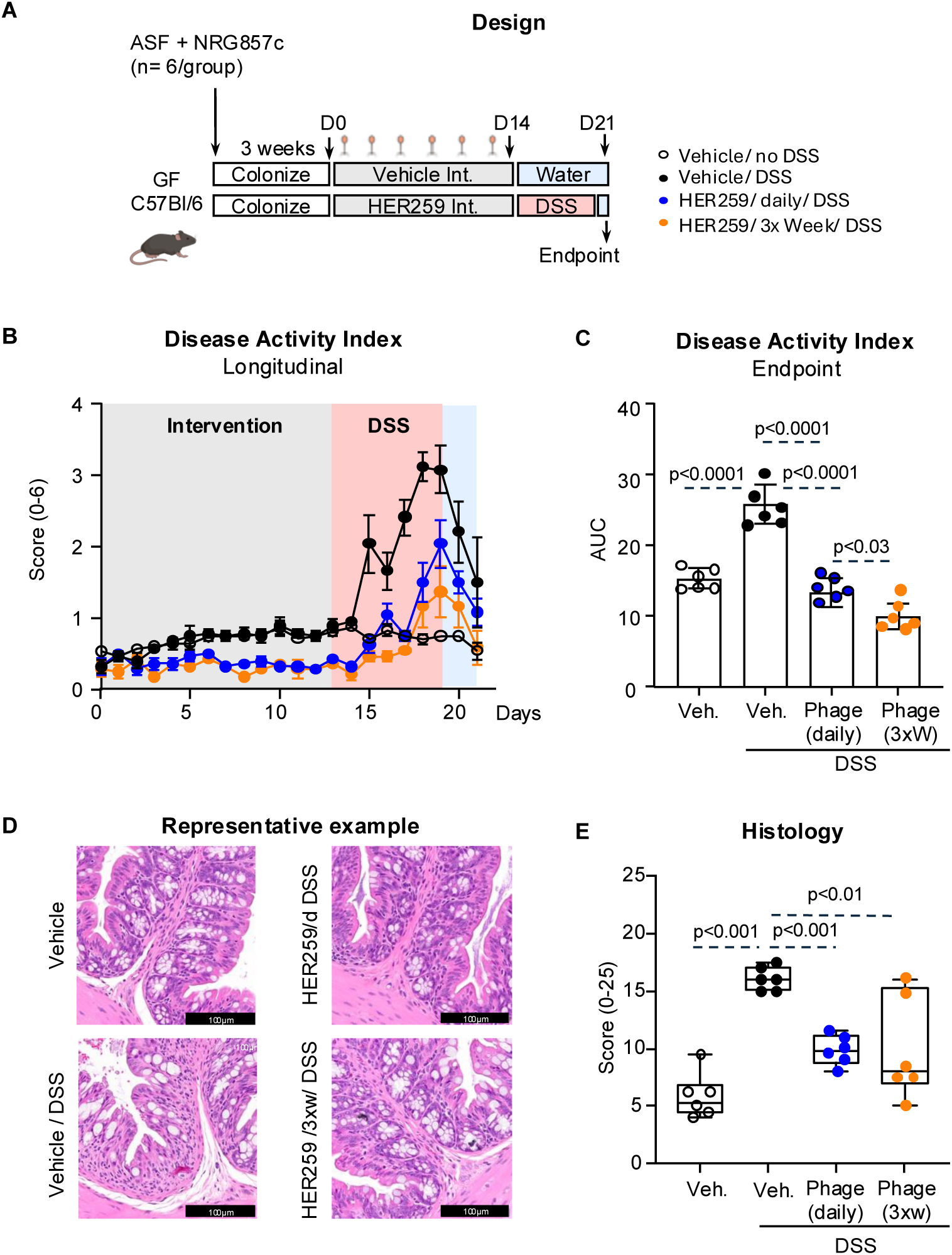
HER259 reduces NRG857c-mediated DSS-induced colitis. (**A**) Germ-free C57Bl/6 mice were co-colonized with ASF and 10^8^ CFU of *E. coli* NRG857c, for 3 weeks. Mice were treated for 2 weeks with either: 1) vehicle daily (PBS with 0.1% bicarbonate); 2) phage HER259 (1×10^9^ PFU/dose; 0.1% bicarbonate) daily; or 3) phage HER259 (1×10^9^ PFU/dose; 0.1% bicarbonate) 3 times/ week. DSS (2%) was administered in drinking water for 5 days followed by 2 days of water. Vehicle treated control mice received water alone. (**B**) Disease activity index (stool consistency + occult blood) analysed longitudinally. (**C**) Area under the curve for disease activity index at endpoint. (**D**) Representative histology examples of cross-sectional proximal colon (40X). (**E**) Histology scores, using a modified grading system (0-25). Statistical significance determined by ANOVA with Tukey post-hoc test.

To investigate if infective viruses are necessary to mitigate colitis, we treated additional mice with heat-inactivated HER259 or an inert protein, bovine serum albumin (BSA) (**Fig S7A**). Neither of these interventions prevented the onset of colitis in ASF+NRG857c mice (**Fig S8B, S8C**). At endpoint, histology scores in mice challenged with heat-inactivated HER259 or BSA were high (**Fig S8D, S8E**). These findings underscore the necessity of live viruses in colitis improvement.

We next evaluated HER259’s capacity to delay spontaneous colitis in *IL-10 ^-/-^* mice under identical colonization conditions (**Fig 4A**). Given the chronic nature of the model, mice were dosed weekly with phage to avert physical distress from continual oral gavage. Weekly phage dosing delayed the onset of NRG857c-mediated colitis in *IL-10 ^-/-^* mice, evidenced by improved stool consistency and decreased incidence of both rectal and occult bleeding (**Fig 4B**). Longitudinal analysis revealed phage-treated mice had reduced disease activity compared with vehicle-treated controls (**Fig 4C**, p<0.01). All vehicle-treated *IL-10^-/-^* mice developed rectal prolapse at endpoint, a clinical sign of colitis activity, while this was absent in phage-treated mice (**Fig 4D**, top). Histological analysis at sacrifice revealed lower inflammation in phage-treated mice compared with vehicle controls (**Fig 4D**, bottom, **Fig 4E**, p<0.05). Thus, live phage administration attenuated the severity of acute and chronic colitis in ASF+NRG857c mice, however the exact mechanisms underlying these improvements required further investigation.

**Fig. 4.**
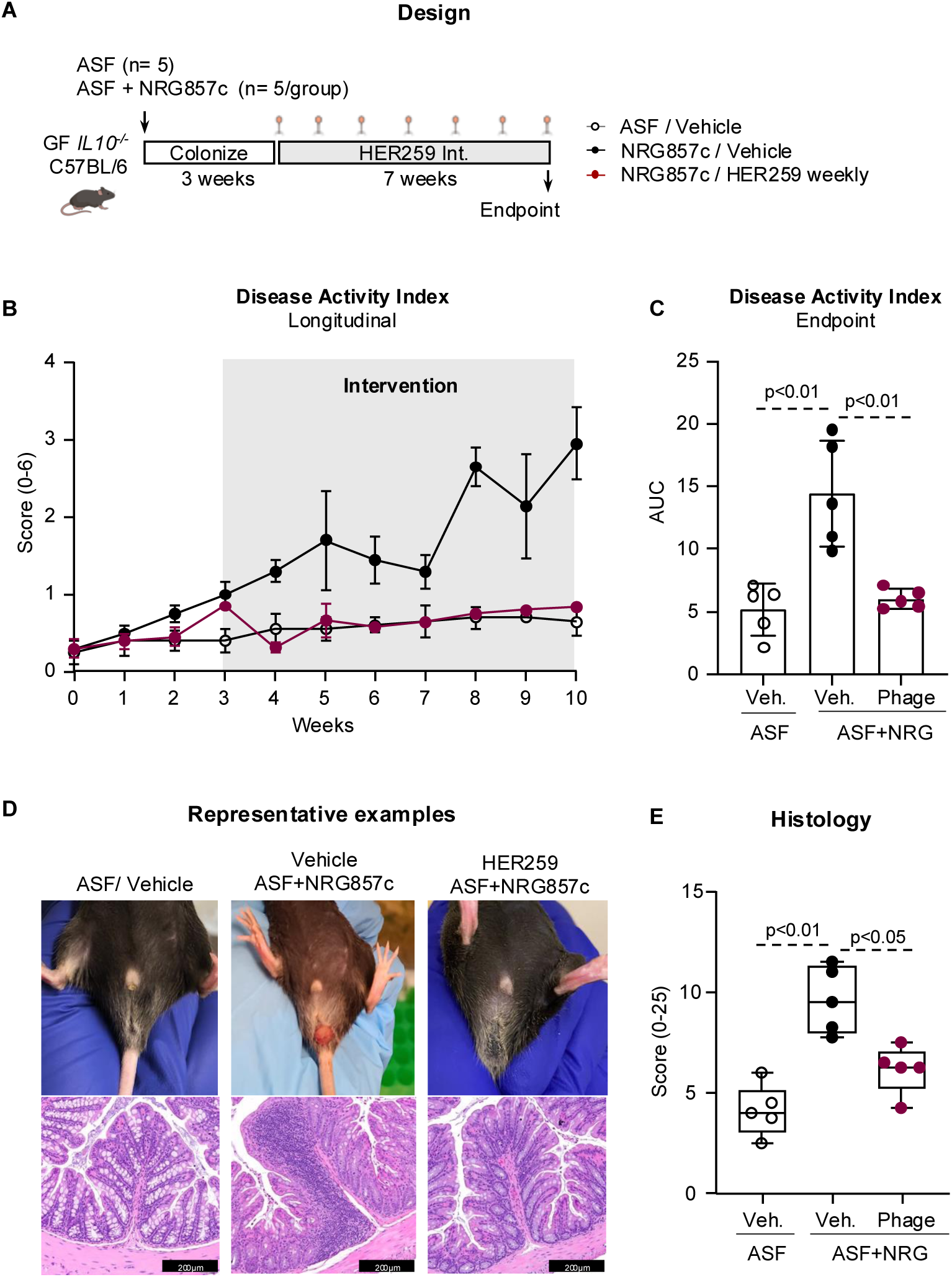
HER259 reduces NRG857c-mediated spontaneous colitis in *Il10*^-/-^ mice. **(A**) Germ-free *Il10*^-/-^ mice were colonized with altered Schaedler flora (ASF) alone, or ASF plus 10^8^ CFU of *E. coli* NRG857c. Three weeks later, mice were treated with either: 1) vehicle daily (PBS with 0.1% bicarbonate); or 2) phage HER259 (1×10^9^ PFU/dose; 0.1% bicarbonate) once per week. Blood in stool and evidence of prolapsed rectum was monitored throughout experiment. (**B**) Disease activity index analyzed longitudinally. (**C**) Area under the curve for disease activity index at endpoint. (**D**) Prolapsed rectums were observed in ASF + NRG857c vehicle treated mice, but not in mice colonized with ASF alone, or phage treated mice (top). Representative histology examples of cross-sectional terminal colon (20X). (**E**) Histology scores, using a modified grading system (0-25). Statistical significance determined by Kruskal-Wallis test with Dunn’s post-hoc test or ANOVA with Tukey post-hoc test.

### HER259 attenuates a key AIEC virulence factor

Readouts for therapeutic success following phage treatment across indications have traditionally focussed on multi-log reductions or complete eradication of the disease-associated bacteria. We thus evaluated the fecal bacterial load of NRG857c and observed a modest, 1-log reduction after 2 weeks of phage treatment, in the absence of colitis, while bacterial load remained stable in vehicle treated mice (**Fig 5A, 5B).** 16S rRNA gene sequencing showed mild non-significant reductions in relative abundance of NRG857c after 2 weeks of intervention, irrespective of phage treatment (**Fig S7A, S7B**) and in vehicle-treated mice (**Fig S7B**), suggesting normal variations of the microbiota over the 5 weeks post colonization. The lack of eradication of NRG857c and any major changes in microbiome composition by phage intervention suggested additional mechanisms for the observed colitis improvement.

**Fig. 5.**
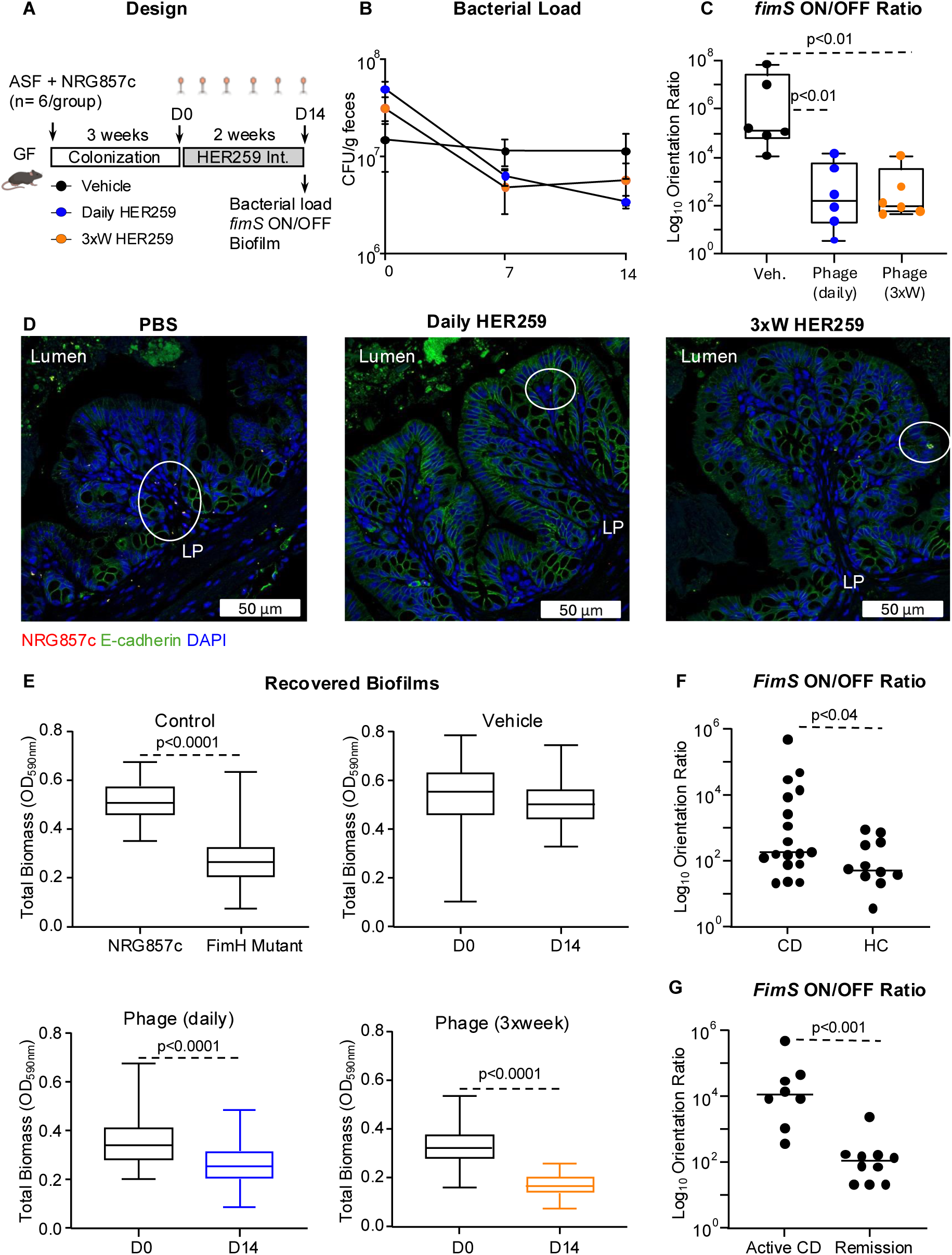
HER259 attenuates key virulence factor of NRG857c. **(A**) Germ-free C57Bl/6 mice were co-colonized with ASF and 10^8^ CFU of *E. coli* NRG857c, for 3 weeks. Mice were treated for 2 weeks with either: 1) vehicle daily (PBS with 0.1% bicarbonate); 2) phage HER259 (1×10^9^ PFU/dose; 0.1% bicarbonate) daily; or 3) phage HER259 (1×10^9^ PFU/dose; 0.1% bicarbonate) 3 times/ week. (**B**) Fecal load of NRG957c during phage intervention. (**C**) Log_10_ Orientation Ratio of *fimS* ON to *fimS* OFF, as measured by qPCR in fecal samples collected at D14. (**D**) Immunostaining in colonic sections of ASF + NRG857c colonized mice treated with vehicle (left) or HER259 (daily, middle; three times/week, right). Yellow represents NRG857c, blue represents DAPI-stained nucleus, and green represents E-cadherin. (**E**) Biofilm forming capacity of recovered NRG857c as assessed using microtiter dish biofilm formation assay. (**F**) Log_10_ Orientation Ratio of *fimS* ON to *fimS* OFF, as measured by qPCR, in CD vs. Health control (HC), and (**G**) Active CD vs. Remission. Statistical significance determined by Kruskal-Wallis test with Dunn’s post-hoc test or ANOVA with Tukey post-hoc test.

Reduced bacterial virulence, including bacterial adhesion capacity, has been reported following phage challenge *in vitro*.(*23–25*) The ability for AIEC to adhere tightly to intestinal epithelial cells, mediated by FimH, is considered one of its central virulence factors exacerbating intestinal colitis.(*26, 27*) Previous reports have shown C57BL/6 mice pre-treated with broad-spectrum antibiotics colonized with AIEC lacking FimH have an attenuated colitis phenotype.(*28*) Thus, we first determined whether FimH is necessary for NRG857c to exacerbate colitis in our gnotobiotic models. For this, we colonized GF mice with ASF+NRG857c Δ*fimH* mutant. Following the microbiota stabilization period, experimental colitis was induced by DSS (**Fig S9A**). Mice colonized with the Δ*fimH* mutant had reduced disease activity (**Fig S9B, S9C** p<0.0001) and lower histological scores compared with ASF+NRG857c wild-type (WT) (**Fig S9D, S9E** p<0.002). Immunohistochemistry revealed that NRG857c Δ*fimH* were restricted to the mucosal surface, suggesting an impaired ability to invade into the lamina propria (**Fig S9F**). (*14*) Endpoint evaluation of cecal contents revealed comparable bacterial loads between mice colonized with NRG857c WT and NRG857c Δ*fimH* (**Fig S9G**). These results supported FimH as a key virulence factor influencing NRG857c’s colitogenic capacity and that its downregulation would yield an attenuated colitis phenotype.

Next, we quantified the relative *fimS* ON/OFF ratio using qRT-PCR,(*29, 30*) for the phase-ON orientation is essential for downstream FimH expression.(*12*) After two weeks of phage intervention, the *fimS* ON/OFF ratio was downregulated 3-fold compared with vehicle-treated mice and irrespective of dosing regimen (**Fig 5C**, p<0.01, respectively). Immunohistochemistry revealed that NRG857c penetrated deep into the lamina propria in vehicle-treated mice (**Fig 5D**, left), while NRG857c in phage-treated mice were restricted to the mucosal surface, irrespective of dosing regimen (**Fig 5D**, middle, right), consistent with observations in NRG857c Δ*fimH* experiments.

To assess whether these attenuated virulence mechanisms were due to HER259 phage binding to NRG857c via FimH, we spot tested HER259 against a NRG857c WT (**Fig S10A**), a NRG857c Δ*fimH* mutant (**Fig S10B**) and a recovered NRG857c WT with *fimS* in the phase-OFF orientation (**Fig S10C**). The observed clearings ruled out that phage infection occurred mainly via FimH.

Another key virulence feature of pro-inflammatory bacteria, including NRG857c, is their predisposition to form biofilms *in vivo*, a mechanism that is, in part, supported by FimH.(*31–33*) Thus, we hypothesized that phage treatment reduced NRG857c’s ability to form biofilms. To test this, NRG857c colonies were isolated from fecal samples before and after phage treatment, prior to colitis induction (**Fig 5A**, D0, D14). Recovered NRG857c isolates were grown overnight in 96-well plates, and the amount of surface-adhered biomass was quantified using crystal violet staining.(*34*) We first compared WT NRG857c against a NRG857c Δ*fimH* mutant (**Fig 5E**; top-left). We observed reduced biofilm formation in wells containing the NRG857c Δ*fimH* mutant compared to wells containing the WT NRG857c (**Fig 5E**, top-left, p<0.0001). There were no differences in the biofilm-forming capacity of recovered WT NRG857c strains from vehicle-treated mice at D0 (pre-treatment) and D14 (post-treatment) (**Fig 5E**, top-right). In contrast, we observed a significant reduction in the biofilm-forming capacity of NRG857c recovered at D14 following daily phage-treatment (**Fig 5E**, bottom-left, p<0.0001) or 3×week phage-treatment (**Fig 5E**, bottom-right, p<0.0001) as compared to WT NRG857c recovered at D0 prior to phage-treatment.

These results collectively support the notion that phage intervention imparts a less virulent bacterial phenotype on surviving NRG857c compared with vehicle-treated counterparts. This is independent of phage-FimH binding and thus unlikely a direct result of phage-receptor loss prior to colitis onset.

### Elevated fimS ON/OFF Ratio present in active Crohn’s disease patients

To validate the clinical relevance of *fimS* regulation as a target for phage outcome in human cohorts, we quantified the *fimS* ON/OFF ratio in feces collected from a subgroup of CD patients and healthy volunteers from the IMAGINE cohort (Table S1). CD patients had elevated *fimS* ON/OFF ratios compared with healthy controls (**Fig 5F**, p<0.04). After stratifying CD patients into those with active flares or in clinical remission, we detected elevated *fimS* ON/OFF ratio in patients with active disease compared to those in remission (**Fig 5G**, p<0.001). The results indicate that known virulence markers in taxa associated with CD(*10, 27*) are increased during active flares, providing a strong rationale that disrupting this mechanism may be key for phage therapy success in clinical translation.

### Sustained administration is required to maintain the clinical effect of HER259

Next, we investigated the longevity of colitis suppression by phage treatment, using GF *IL-10^-/-^* mice colonized with ASF, with or without NRG857c. Mice were treated weekly with HER259 or vehicle for seven weeks, and then monitored for 6 more weeks following cessation of phage treatment (**Fig 6A**). Consistent with data in Fig 4, phage intervention delayed the onset of colitis, until week 14, when phage-treated mice began displaying disease signs, including reduced stool consistency and increased rectal bleeding, as illustrated by their combined DAI scores (**Fig 6B, 6C)**. No differences in DAI were observed by week 16 between phage- and vehicle-treated mice (**Fig 6B)**, despite there being a two-log difference in the NRG857c bacterial load (**Fig 6D**). Histological analysis at sacrifice further revealed no differences between the two treatment groups (**Fig 6E, 6F)**, suggesting a reversion of NRG857c to a pro-inflammatory phenotype. To test this, we quantified the *fimS* ON/OFF ratio at week 3 (pre-phage treatment), week 10 (post-phage treatment), and at week 16 (endpoint) (**Fig 6A**). There were no differences in the orientation ratio between intervention groups at week 3 **(Fig 6G**, left), while at week 10, there was a 2-fold lower *fimS* ON/OFF ratio in phage-treated mice compared to vehicle-treated mice (**Fig 6G**, middle, p<0.01). At week 16, we observed no differences in the *fimS* ON/OFF between the two intervention groups (**Fig 6G**, right), indicating that surviving NRG857c in phage-treated mice reverted to a pro-inflammatory phenotype following cessation of the phage treatment. This reversion coincided with an increase in clinical scores of colitis in phage-treated mice following cessation of the treatment, a non-significant increase in the relative abundance of NRG857c in HER259-treated mice (**Fig S11B**), and no shift in beta diversity following cessation of HER259 treatment (**Fig S11D**). Thus, these results support that phage treatment is necessary to suppress colitis through downregulation of *fimS* ON/OFF orientation ratio, and that chronic phage administration is required to maintain is beneficial effect.

**Fig. 6.**
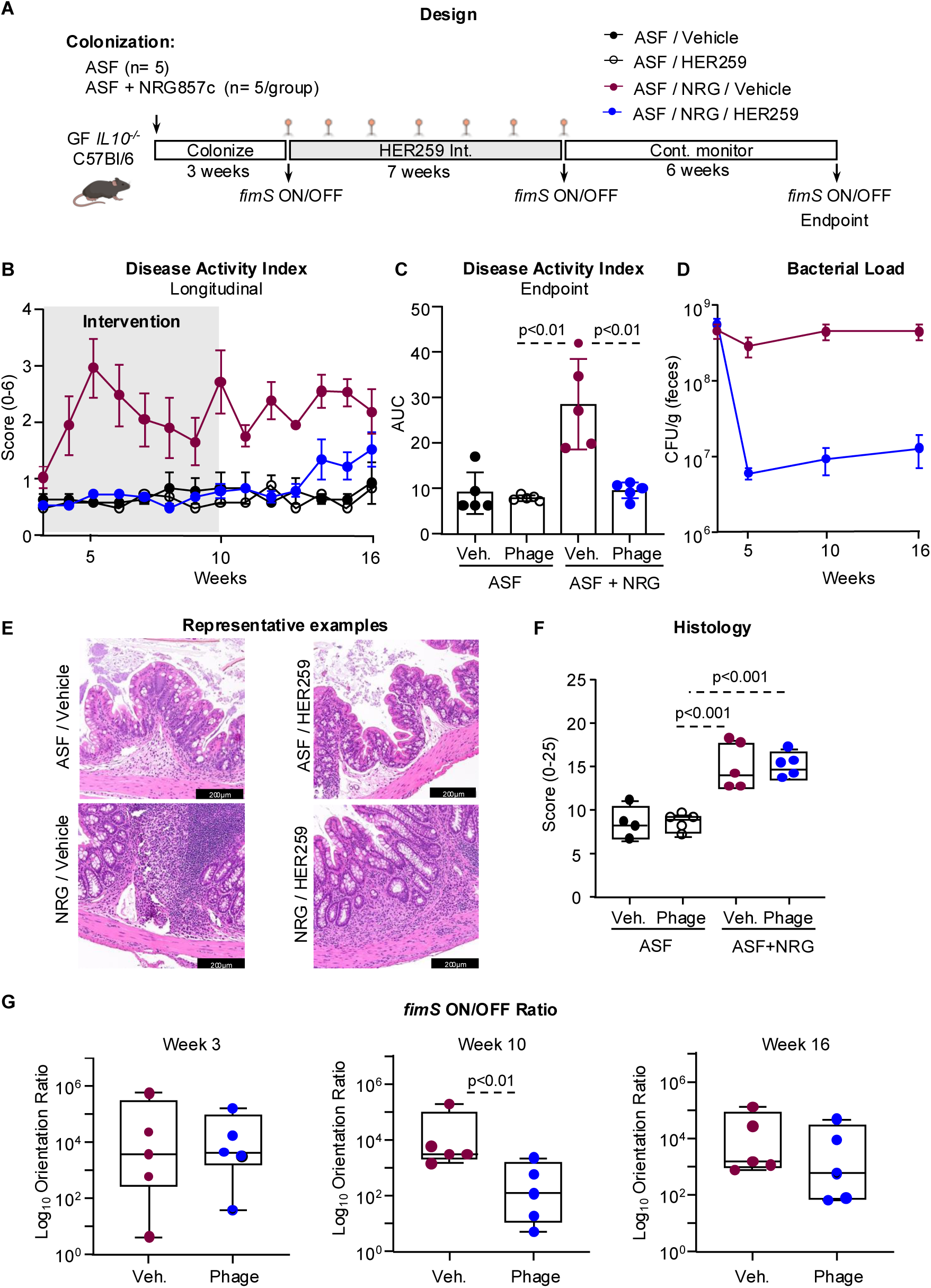
Colitis phenotype returns following cessation of HER259 intervention. (**A**) Germ-free *Il10*^-/-^ mice were colonized with ASF alone, or ASF plus 10^8^ CFU of *E. coli* NRG857c. Three weeks later, mice were treated with either: 1) vehicle daily (PBS with 0.1% bicarbonate); or 2) phage HER259 (1×10^9^ PFU/dose; 0.1% bicarbonate) once per week. Stool consistency and occult blood were monitored weekly throughout experiment. (**B**) Disease activity index analysed longitudinally. (**C**) Area under the curve for disease activity index at endpoint. (**D**) Bacterial loads of ASF-NRG857c-colonized mice treated with Vehicle or phage. (**E**) Representative histology examples of cross-sectional proximal colon (20X). (**F**) Histological scores using a modified grading system (0-25). (**G**) Log_10_ Orientation Ratio of *fimS* ON to *fimS* OFF, as measured by qPCR, in mice treated with Vehicle or phage at Week 3 (left), Week 10 (middle), and Week 16 (right). Statistical significance determined by Kruskal-Wallis test with Dunn’s post-hoc test or ANOVA with Tukey post-hoc test.

### HER259 prevents colitis reactivation

To mimic the relapsing-remitting cycle of inflammation observed in patients with IBD, we tested HER259’s therapeutic potential in a colitis reactivation protocol. GF C57BL/6 mice were colonized with ASF with or without NRG857c and then exposed to a first cycle of low-dose DSS, followed by phage 3xweek (or vehicle) and a second DSS cycle (**Fig 7A**). Analysis over the 2 DSS cycles demonstrated progressive increase in colitis severity in ASF+NRG857c receiving vehicle **(Fig 7B)**. At endpoint vehicle-treated ASF+NRG857c mice had higher colitis severity than ASF alone (p<0.0001), and this was attenuated by phage (**Fig 7C**, p<0.0001). Phage also decreased colonic histology scores in ASF+NRG857c mice as compared to vehicle-treated mice (**Fig 7D, 7E** p=0.03). To further analyse differences in disease activity progression, we stratified groups into the distinct treatment periods (**Fig S12)**. ASF mice experienced an increase in DAI scores from first to second DSS cycle, irrespective of treatment with phage or vehicle (**Fig S12C, S12D,** p<0.0001 and p<0.0001, respectively). While vehicle-treated ASF+NRG857c experienced a 160% increase **(Fig S12E,** p<0.0001), phage-treated ASF+NRG857c showed a 10% decrease in DAI between the first and second DSS cycle, supporting phage’s ability to prevent exacerbation of inflammation after a second bout of colitis (**Fig S12F**).

**Fig. 7.**
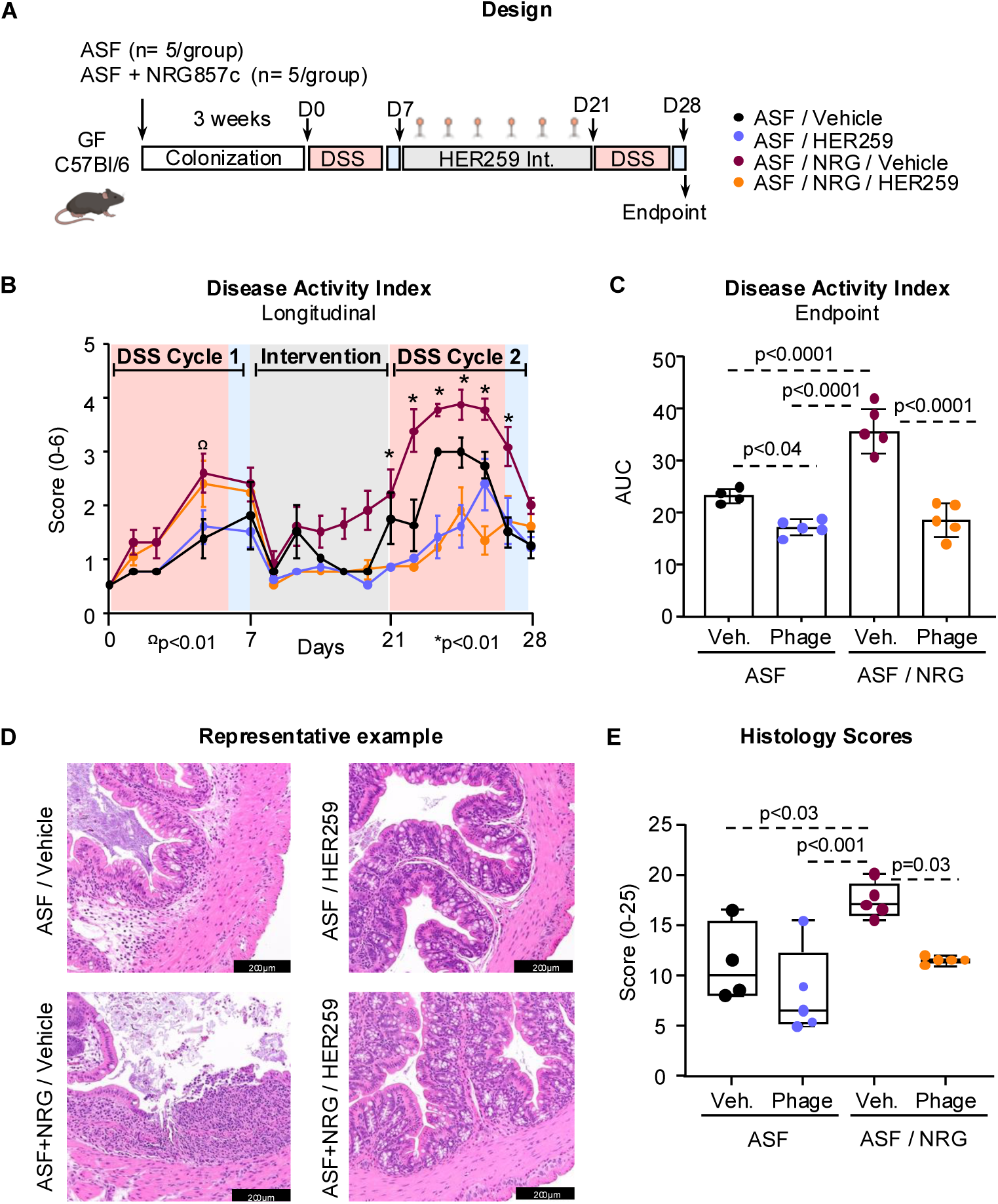
HER259 treatment prevents colitis reactivation. (**A**) Germ-free C57Bl/6 mice were co-colonized with ASF and 10^8^ CFU of *E. coli* NRG857c, for 3 weeks. DSS (2%) was administered in drinking water for 5 days followed by 2 days of water to induce inflammation (DSS cycle 1). Following the first DSS cycle, mice were treated for 2 weeks with either: 1) vehicle daily (PBS with 0.1% bicarbonate); or 2) phage HER259 (1×10^9^ PFU/dose; 0.1% bicarbonate) 3 times/ week. DSS (2%) was administered in drinking water for 5 days followed by 2 days of water (DSS cycle 2). Mice were monitored for stool consistency, and presence of occult blood. (**B**) Disease activity index analysed longitudinally (^Ω^p<0.01 ASF vs. ASF+NRG857c; *p<0.01 ASF+NRG857c/Vehicle vs. ASF+NRG857c/HER259). (**C**) Area under the curve for disease activity index at endpoint. (**D**) Representative endpoint histology of cross-sectional proximal colon taken at 20X. (**E**) Colonic histological scores using a modified grading system (0-25). Statistical significance determined by Kruskal-Wallis test with Dunn’s post-hoc test or ANOVA with Tukey post-hoc test.

### HER259 boosts the effects of a sub-therapeutic dose of Budesonide

IBD therapies, such as corticosteroids, other immunosuppressors, and biologics are usually combined to overcome failures observed in patients with IBD.(*35*) Thus, we tested the efficacy of HER259 in combination with budesonide, a corticosteroid medication delivered orally or by enema, and used to treat mild-moderate colitis.(*36*) We colonized GF C57BL/6 mice with ASF and NRG857c, and treated them with phage 3xweek or vehicle, before inducing colitis by DSS (**Fig 8A**). Midway through the DSS cycle, mice received a sub-therapeutic dose of budesonide(*37*) for 5 days (**Fig 8A**). Combined phage-budesonide treatment reduced DAI scores compared with budesonide- or phage-treatment alone (**Fig 8B**). Endpoint analysis of DAI scores revealed that combined phage-budesonide treatment resulted in better clinical outcomes compared with individual treatments (**Fig 8C**, p<0.001, p<0.05 respectively). Mice with combined treatment had significantly reduced histological scores at endpoint compared with vehicle-alone (**Fig 8D, 8E**, p<0.0001), budesonide-alone (**Fig 8D, 8E**, p<0.05), or phage-alone treatment (**Fig 8D, 8E**, p<0.05). Thus, prophylactic phage treatment boosts the response to a subtherapeutic dose of budesonide.

**Fig. 8.**
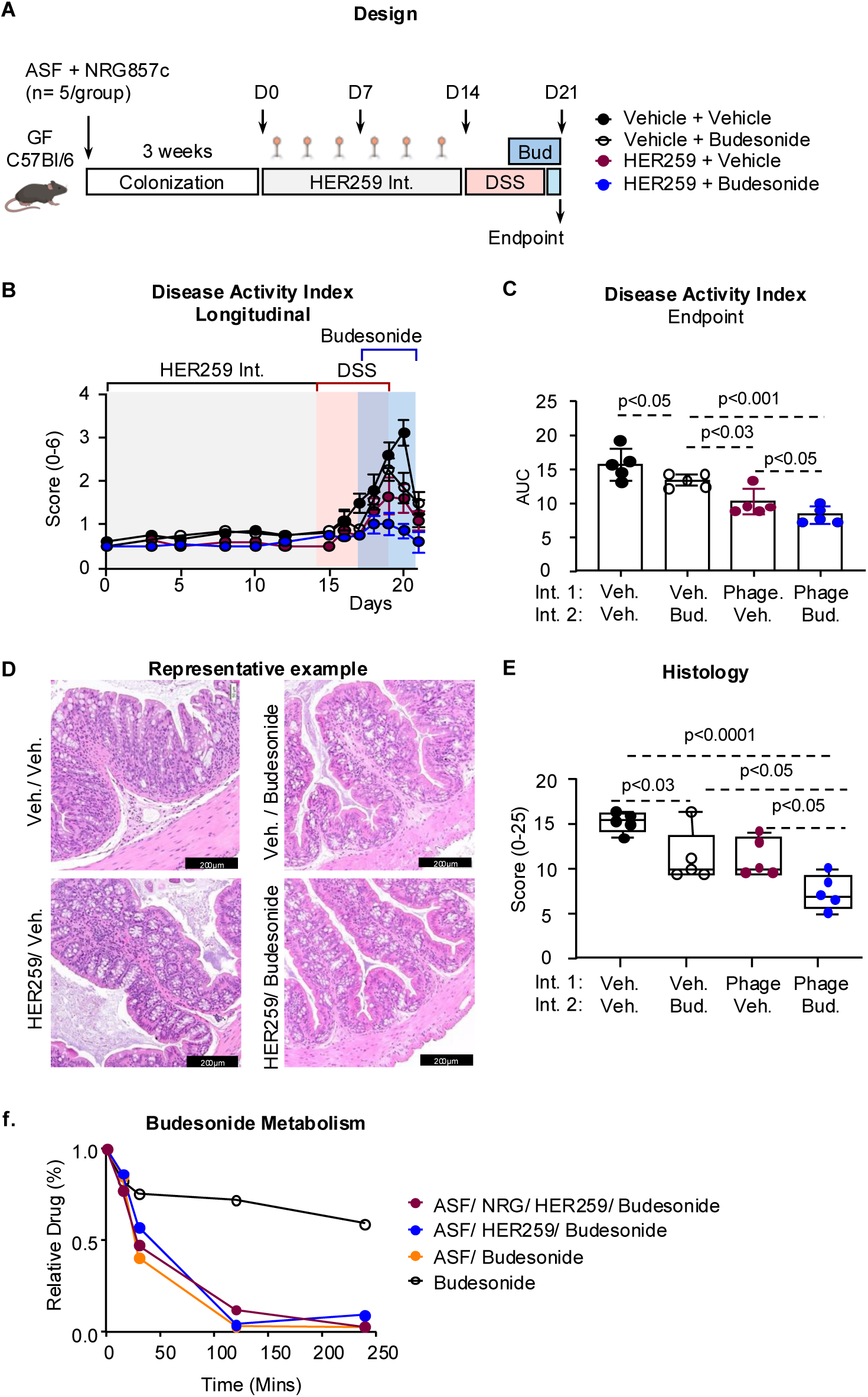
Phage intervention enhances host response to corticosteroid. **(A**) Germ-free (GF) C57Bl/6 mice were colonized with ASF plus 10^8^ CFU of *E. coli* NRG857c. Three weeks later, mice were treated with either: 1) vehicle daily or 2) phage HER259 (1×10^9^ PFU/dose) daily for 14 days. Mice then received DSS (2%) in drinking water for 5 days followed by 2 days of water. A low dose of budesonide (Bud; 2 ug/daily by oral gavage) was initiated halfway through DSS administration and continued to endpoint. (**B**) Disease activity index analysed longitudinally. (**C**) Area under the curve for disease activity index at endpoint. (**D**) Representative histology examples of cross-sectional proximal colon (20X). (**E**) Colonic histological scores using a modified grading system (0-25). (**F**) Residual active budesonide as determined by HPLC analysis. Statistical significance determined by Kruskal-Wallis test with Dunn’s post-hoc test or ANOVA with Tukey post-hoc test.

### Therapeutic effect of Budesonide is independent of microbial drug metabolism

Phage treatment has previously been suggested to alter metabolic functions of the gut microbiome in gnotobiotic mice.(*38*) To this end, we hypothesized that HER259 might alter the gut microbiome’s ability to metabolize budesonide, given this drug is administered luminally.(*39*) Thus, we anaerobically incubated budesonide with fecal slurries containing ASF, NRG857c, and phage, and then quantified residual budesonide metabolites using HPLC at regular intervals. No differences were observed in the residual metabolites following phage challenge across experimental conditions (**Fig 8F**), suggesting the gut microbiota likely does not affect budesonide metabolism or increase its availability in the model.

## Discussion

IBD is a complex condition in which microbial factors are thought to induce and drive inflammation in genetically susceptible hosts.(*1–3*) Given the non-specific nature and adverse events associated with antibiotics, interest in phage therapy is growing, with the expectation that their clinical efficacy will result from bacterial load reduction. We show here that phage reduces bacterial load but also attenuates bacterially mediated colitis by downregulating virulence markers of *E. coli* NRG857c. The attenuated bacterial and colitis phenotypes were ubiquitous across animal models, promoting enhanced clinical responses to subtherapeutic doses of the corticosteroid, budesonide, following phage administration. We identified a previously unknown mechanism in which phage HER259 downregulates proinflammatory behavior of a CD-associated isolate, mitigating colitis and improving host responses to budesonide.

To date, clinical trials evaluating phage therapy in IBD have used cocktails designed to reduce bacterial load, but this approach overlooks the complex nature of IBD pathogenesis. To study some of these interactions, we developed novel gnotobiotic mouse models colonized with a defined (ASF-like) community combined with, or without, known pro-inflammatory *E. coli* strains, such as NRG857c, isolated from patients with CD. Our models differed from their predecessors in which mono-colonization or pre-treatment with antibiotics followed by AIEC colonization were employed.(*20, 40*) We began by assessing the colitogenic capacity of three clinically relevant AIEC-like strains: C0004, a urinary isolate, and two CD isolates, LF82 and NRG857c. The latter, NRG857c, demonstrating the greatest capacity to exacerbate colitis. Based on robust pre-clinical *in vitro* assays, including kill curves and biofilm challenges, we identified HER259 as a novel phage candidate for these strains. When translated to *in vivo* testing, our targeted phage interventions resulted in modest NRG857c load reductions in feces, lower in magnitude from previous reports using other phages, which may be attributed to the microbial stability of ASF microbiome in this study as compared with mono-colonization systems, or perturbation of background microbiome by antibiotic pretreatments.(*20, 40*) Because the target bacterium was not eradicated and only modest load reductions were observed, we posit that other mechanisms are at play for the marked improvement in colitis by phage therapy.

Virulence of AIEC is characterized by their ability to invade and tightly adhere to the mucosa, a process that is in part mediated by FimH. Phages are known to exert selective pressures that drive phenotypic and genotypic adaptations, often leading to reduced bacterial virulence.(*23–25*) In accordance with this, we observed a 2-fold reduction in the *fimS* ON/OFF ratio, suggesting reduced FimH expression following phage intervention.(*12, 41*) Immunohistochemistry revealed reduced bacterial invasion into the lamina propria following phage treatment, a feature that was likewise observed in mice colonized with NRG857c *ΔfimH*. Moreover, recovered NRG857c isolates from phage-treated mice had lower biofilm-forming capacity *in vitro*, which is known to be mediated, at least in part, by FimH.(*33*) These results indicate that HER259 intervention imparts an attenuated virulence phenotype on surviving NRG857c. While novel in the context of IBD, attenuated bacterial virulence following phage therapy has been reported when treating multidrug resistant bacterial infections.(*42*) Moreover, lytic phages have been shown to induce bacterial phase variations in the gut, suggesting invertible promotor region plasticity during active flares could drive attenuated clinical phenotypes.(*43, 44*) To investigate whether elevated adhesion virulence factors have potential as targetable mechanism in humans, we procured samples from a cross-sectional cohort of patients with IBD. We quantified fecal *fimS* ON/OFF ratios and determined an increase in patients with active CD, versus those in remission, and healthy controls. Collectively, our findings have potential to optimize clinical trial design for evaluating phage therapy in IBD by improving patient selection based on high fecal *fimS* ON/OFF ratio and the incorporation of attenuation of virulence as a readout.

To replicate the relapsing-remitting inflammatory cycles characteristic of IBD, we assessed the therapeutic efficacy of HER259 using a newly established colitis reactivation model. Following an initial episode of DSS-induced colitis, targeted phage intervention attenuated disease reactivation during a second DSS challenge. These results reinforce both the preventive and therapeutic potential of using phage therapy in the context of IBD, especially under more clinically relevant conditions.

Existing IBD drugs are often used in combination, given the complexity and multifactorial origin of the disease,(*35*) and we postulated phage therapy for IBD would be no different. Our experiments demonstrate that host responsiveness to a sub-therapeutic dose of budesonide is enhanced when preceded by phage intervention. This effect was independent of microbial-based metabolism of the drug(*45*) as HPLC analysis revealed no changes in active budesonide levels following phage challenge. Our results support the hypothesis that improved response to budesonide is due to the general attenuation of NRG857c virulence and its lower inflammatory potential. This is clinically relevant, given that half of IBD patients experience a secondary loss of response to pharmacological therapies within 12 months.(*46*) While underlying mechanisms for treatment failure remain unknown, it has been proposed that some of these flares are caused by the presence of pro-inflammatory bacteria, including AIEC-like strains.(*10, 27, 47–50*) Phages have long demonstrated synergistic effects with antibiotics in combating multi-drug-resistant infections. This synergy has been extensively studied both pre-clinically and clinically, establishing a widely held belief that phages are primarily useful in this context.(*51*) Our results align with these findings suggesting phage therapy in IBD could synergize with existing therapies.

The results presented here are promising for the future of precision therapeutics, however there are some limitations to this study. First, we selected HER259 and NRG857c as a model phage-bacterial pairing. While NRG857c may not be present in the microbiota of all patients with IBD, it provided a reductionist framework to evaluate mechanistic new knowledge in this area. Importantly, our approach leveraged a clinically relevant AIEC. (*18, 52*) Second, we used a simplified ASF-like microbiome which we acknowledge does not fully capture the diversity of the human gut. However, the model allowed us to discriminate phage effects towards the single *E. coli* that was target, as no other Enterobacteriaceae was present. Finally, host immune responses potentially induced by phage(*53, 54*) were not investigated. Elucidating these mechanisms, which are beyond the scope of this study, will be essential for advancing phage therapy applications in the gut to treat IBD.

In summary, using phage HER259 and NRG857c as a model phage-proinflammatory bacterium pair, we reveal a new mechanism by which targeted phage therapy disrupted key virulence markers in the CD-associated *E. coli* strain, reducing colitis, preventing reactivation, and enhancing therapeutic responses to budesonide. By reducing bacterial virulence mechanisms without disrupting microbial balance, phage-based treatments align with personalized microbial therapeutics. Our study offers potential to improve the future design of phage intervention trials in IBD, by selecting patients that would most benefit from virulence attenuation and testing synergies with existing conventional treatments to prevent dose escalation, treatment failure, and reduce adverse events.

## Materials & Methods

### Bacterial and phage culture preparation

Pre-cultures of the bacterial host were prepared by inoculating BD Bacto™ Tryptic Soy Broth (TSB; DF0370-17-3, Fisher Scientific) from glycerol stocks and incubating overnight at 37 °C with shaking. The resulting culture was diluted 1:100 into 250 mL of TSB broth, followed by the addition of 10 μL of HER259 phage stock (10^12^ plaque forming units (PFU)/mL). Phage-infected cultures were incubated at 37 °C with shaking for 5 hours. After incubation, the culture was centrifuged at 7,000×g for 15 minutes to pellet the bacteria. The phage-containing supernatant was collected and stored at 4 °C.

### HER259 dose preparation

HER259 was propagated as previously described above using *E. coli* NRG857c. HER259 doses were prepared by polyethylene glycol (PEG)-purification. Briefly, a mixture of 20% (w/v) PEG solution and 2.5M NaCl solution was added to the crude phage lysate with a volumetric ratio of 1:6 and incubated at 4℃ overnight. Phage was pelleted by centrifugation (5000×g, 45 min, 4℃). The pellet was resuspended in 10 mL of PBS and incubated at 4℃ for 2 hrs. This purification step was repeated twice. PEG-purified HER259 solutions were sterile filtered using a 0.2μm filter. HER259 titers were determined by double layer agar overlay method and adjusted to 1×10^9^ PFU/mL.

### Mice

C57BL/6 WT and C57BL/6NTac-Il10^em8Tac^ (*IL-10^-/-^)* mice germ-free mice were generated by 2-stage embryo transfer at McMaster’s Axenic Gnotobiotic Unit (AGU) and were bred under germ-free conditions in flexible film gnotobiotic isolators. Male and female mice were used in all experiments. All mouse experiments were approved by the McMaster University Animal Care Committee and McMaster Animal Research Ethics Board in an amendment to the Animal Utilization Protocol (# 210930).

### Mouse colonization

Colonizer mice containing the Altered Schaedler-like Flora (ASF) were sacrificed and their cecal contents collected into a sterile tubed of PBS containing L-cysteine (0.1% w/v). 100 μL of the homogenized cecal contents were then orally gavaged into a germ-free mouse.

Immediately thereafter, 100 μL of *E. coli* was orally gavage into the mice. Microbiota of newly colonized mice was allowed to stabilize for three weeks.

### Colitis models and evaluation

Colitis severity was investigated in 9 to 14-week-old germ-free C57BL/6 and C57BL/6NTac-Il10^em8Tac^ (*IL-10^-/-^)* colonized with ASF with or without *E. coli*.

Acute colitis was induced by 2% dextran sodium sulfate (DSS, MW 36–50 kDa; MP Biomedicals) in drinking water followed by two days on water only. DSS intake and general health condition were monitored daily. *IL-10^-/-^* colonized mice experienced spontaneous colitis development and did not receive DSS.

Disease Activity Index (DAI, score 0-6) was a composition of stool consistency and occult blood as previously described.(*55*) DAI was evaluated in both C57BL/6, DSS-induced, and *IL-10^-/-^* colitis mouse models.

### Histological analysis of DSS-induced & IL-10^-/-^ colitis models

Colonic tissues were fixed using 10% formalin solution and embedded in paraffin. Colonic sections (5 µm) were stained with hematoxylin and eosin (H&E) and examined under light microscopy. The degree of intestinal inflammation was blindly evaluated using the established cooper scoring method (score 0-4),(*56*) and a modified pathology score (score 0-25).(*55*) Images were acquired using ImagePro Plus (Media Cybernetics, Rockville, USA).

### Detection and quantification of fimS ON/OFF ratio by qPCR

Quantification of the orientation of the *fimS* switch was performed by quantitative PCR (qPCR) with SsoFast EvaGreen Supermix (1725201, Bio-Rad) according to the manufacturer’s recommendations. qPCR was carried out with 2 ng of genomic DNA extracted from mouse fecal samples using a custom protocol as previously described.(*57*) Primers CMD1246 and CMD1248 were used to amplify the ON orientation, while the CMD1247 and CMD1248 primers amplified the OFF orientation (Table S2).(*58*) The C_T_ of the on and off orientations was normalized to the C_T_ of the *E. coli* 16S gene region. (*59*) Fold change was calculated using the standard curve method.(*60*)

### Immunohistochemistry

Slides from formalin-fixed tissue samples were deparaffinized and subjected to antigen retrieval by steaming for 30 minutes in a sodium citrate buffer (10mM sodium citrate, pH 6.0, 0.05% Tween 20). Sections were permeabilized with 0.2% Triton X-100. Endogenous mouse IgG was blocked using the Mouse on Mouse (M.O.M.) Immunodetection Kit (BMK-2202; Vector) according to manufacturer’s specifications. Sections were blocked (hanks balanced salt solution, 2% goat serum, 2% fetal bovine serum, 0.4% Triton X-100) and incubated with antibodies against E. coli O83 (85077; SSI Diagnostica; diluted 1:200) and E-cadherin (CM1681; ECM Biosciences; diluted 1:200) overnight. Sections were then incubated with goat anti-rabbit Alexa Fluor 568 (Invitrogen; 1:200) and goat anti-mouse Alexa Fluor 488 (Invitrogen; 1:2000) for 1 hour. Hoechst 33342, trihydrochloride trihydrate (H3570; ThermoFisher Scientific) was applied for 15 minutes prior to mounting coverslips with ProLong Diamond Antifade (P36965; ThermoFisher Scientific). Images were taken at 60x with oil immersion using a Nikon A1R Inverted confocal microscope.

### Patient samples

Fecal samples from 18 patients with CD (8 active, 10 remission) and 11 healthy controls were obtained from participants of the Inflammation, Microbiome, and Alimentation: Gastrointestinal and Neuropsychiatric Effects (IMAGINE) cohort.(*61*) Disease classification was determined by self-reported patient survey and confirmed using IDK Calprotectin ELISA (MRP8/14, Immundiagnostik AG). See Table S1 for demographic details.

### Budesonide preparation and administration

Budesonide was prepared aseptically at 2 µg/day in 1% Dimethyl sulfoxide (DMSO) /PBS. Germ-free 9–14-week-old C57BL/6 mice, colonized with ASF-like communities with or without E. coli, were treated with HER259 (1×10⁹ PFU/dose) three times per week for two weeks. After HER259 treatment, mice received 2% (w/v) DSS in drinking water for five days, followed by two days of water. Beginning on day two of DSS, mice were dosed daily with budesonide or vehicle until endpoint. Mice were monitored daily for general health and colitis indicators (stool consistency, occult blood), and colitis severity was assessed via histology.

### Statistical Analysis

Data analysis was conducted using GraphPad Prism 9 (GraphPad Software, USA). For comparisons involving more than two groups, one-way analysis of variance (ANOVA) followed by Tukey’s multiple comparisons test was employed. Microbiota data were transformed (logarithmic, square root, reciprocal, or inverse log) as necessary to satisfy assumptions of normality. For comparisons between two groups, either an unpaired Student’s t-test or the non-parametric Wilcoxon rank-sum test was applied, depending on data distribution. For analyses involving more than two non-normally distributed groups, Kruskal–Wallis tests were performed, followed by Dunn’s post hoc test with Holm correction, where appropriate.

## Acknowledgements

KJ holds a Vanier Canada Graduate Scholarship from the Canadian Institute for Health Research. Z.H. holds a Canada Research Chair (Tier 2) in Bacteriophage Bioengineering. EFV holds a Canada Research Chair (Tier 1) in Microbial Therapeutics and Nutrition in Gastroenterology. The authors thank the staff at the Axenic Gnotobiotic Unit and Genomics facility at McMaster University. We also thank the patients that donated samples for this study.

## Author contributions

K.J. conceived, designed, and executed the experiments, analyzed the data, prepared the figures, and wrote the initial draft of the manuscript. H.G. and A.H. made important contributions to *in vivo* studies. M.C. performed 16S rRNA gene sequencing analysis. M.B.J., A.F., H.H., J.W., made important contributions to *in vitro* studies. M.Z. and C.S. supported preparation and analysis of immunohistochemistry. P.M. supplied IMAGINE patient samples. M.S., P.B., provided scientific editing, and feedback on data analysis. Z.H. and E.F.V. conceptualized and supervised the project, and guided the experimental design, data analysis and manuscript writing.

## Data and materials availability

All data associated with this study are present in the paper or the Supplementary Materials. Information about all commercially available reagents is provided in Materials and Methods and table S2.

