## Supplementary Materials for "Phage intervention improves colitis and response to corticosteroids by attenuating virulence of Crohn’s disease-associated bacteria"

#### Supplementary Methods

*HER259 serial passaging*

To increase the infectivity of HER259 against NRG857c, we serially passaged the phage as previously described. *(62, 63)* HER259 was prepared as described above. Filtered supernatants were collected and cultured in fresh preculture solutions. This process was repeated for 10 cycles.

*Combined Disease Activity Index*

Disease Activity Index (DAI, score 0-6)*(3)* was a composition of stool consistency (firm, score = 0; mild diarrhea = 1; moderate diarrhea = 2; severe diarrhea = 3) and occult blood (absent, score = 0; positive test, score = 1; visible blood, score = 2; free flowing blood, score = 3).

*Histology analysis following phage HER259 intervention*

We used the following parameters as previously described.*(64)* a) Presence of PMN and erythrocytes in luminal exudate (absent, score = 0, scant = 1, moderate = 2, dense = 3); b) Epithelial damage and desquamation (no pathological change, score = 0, mild regenerative change = 1, moderate with patchy desquamation = 2, severe with diffuse desquamation = 3); c) Acute inflammatory infiltrate of the mucosa (PMNs) (absent, score = 0, scant = 1, moderate = 2, dense = 3); d) Mononuclear cell infiltrate of the mucosa (absent, score = 0, one small aggregate = 1, more than one aggregate = 2, several aggregates = 3); e) Goblet cell depletion (abundant number of goblet cells, score = 0, mild depletion = 1, moderate = 2, severe = 3); f) Cryptitis and crypt abscesses (absent, score = 0, isolated cryptitis = 1, diffuse cryptitis = 2, crypt abscesses = 3); g) Architectural damage (no pathological change, score = 0, mild and isolated architectural change = 1, moderate change = 2, severe and diffuse changes = 3); h) Edema (absent, score = 0, mild edema = 1, moderate = 2, severe = 3). An average of all parameters was used to determine the microscopic inflammatory score (0–3).

*Biofilm model*

Biofilms were prepared aerobically in sterile 96-well plates (10861-564, VWR). LF82 and NRG857c pre-cultures were grown overnight at 37 °C with shaking. 1:10 Subcultures were prepared in fresh TSB and incubated at 37 °C until $OD_{600nm}$ reached 0.6. Biofilms were seeded by adding 200 µL of subculture or TSB (control) to 96-well plates and incubated at 37 °C, 180 RPM for 24 h. For untreated biofilm assays, media were removed, wells washed 3× with phosphate buffered saline (PBS; BP2944100, Fisher Scientific), stained with 0.05% crystal violet (CV) for 20 min, washed, and dried overnight. The next day, biomass was dissolved in 30% acetic acid and absorbance measured at OD590nm (BioTek plate reader). For treated biofilm assays, 200 µL of treatment solution (PBS or phage) was added post-seeding and incubated for 24 h. Wells were then processed as above (PBS wash, 0.05% CV stain, drying, and acetic acid solubilization) before $OD_{590nm}$ measurement (BioTek plate reader).

*16S ribosomal RNA gene sequencing*

Bacterial genomic DNA was extracted as previously described.*(65)* The variable regions 3-4 of the 16S rRNA gene was amplified and sequenced using the MiSeq Illumina platform via an established protocol.*(66)*

Sequences were processed in R (version 4.4.2) using the package Divisive Amplicon Denoising Algorithm 2 (DADA2)*(67)* and the SILVA reference database (version 138.1).*(68)*

Phylogenetic tree of sequences were calculated using FastTree 2*(69)* and data were explored using the phyloseq package.*(70)* Beta-Diversity was calculated using the Aitchison distance. Permutational Multivariate Analysis of Variance (PERMANOVA) was used to calculate differences between groups.*(71)* Taxonomic differences were evaluated using a generalized linear model with a negative binomial distribution.*(72)* Statistical differences between groups were assessed using estimated marginal means via the emmeans package.(*73)*

Data available at https://www.ncbi.nlm.nih.gov/bioproject BioProject ID: PRJNA1272099

*Budesonide stability & high-performance liquid chromatography analysis*

Budesonide stability was assessed using high-performance liquid chromatography HPLC analysis as previously described.*(74)* Fecal slurries of ASF and budesonide (1078201, Sigma) were prepared and incubated anaerobically with NRG857c ($10^8$ colony forming units (CFU) / mL) and HER259 ($10^9$ PFU / mL). Aliquots were sampled at regular time intervals (15 mins, 30 mins, 120 mins, 240 mins) and centrifuged at 5000 x g. Supernatants were collected, and ice chilled acetonitrile (34851, Sigma) was added to each sample in a 1:1 ratio. Samples were analyzed by injecting 5 µL into an Agilent 1290 Infinity II HPLC system equipped with a diode array detector (DAD) (Agilent, Santa Clara, CA, USA) set to 254 nm. Budesonide was separated on an Agilent (Santa Clara, CA, USA) Eclipse Plus Phenyl-Hexyl (100 mm x 2.1 mm, i.d., 1.8 µm) analytical column using mobile phases consisting of (A) water and (B) acetonitrile and a constant flow of 0.3 mL/min. The autosampler was maintained at 10˚C throughout the analysis, and the analytical column was maintained at 40 ˚C. The separation was achieved by a linear gradient over a run time of 22 minutes.

### Supplementary Figures & Tables

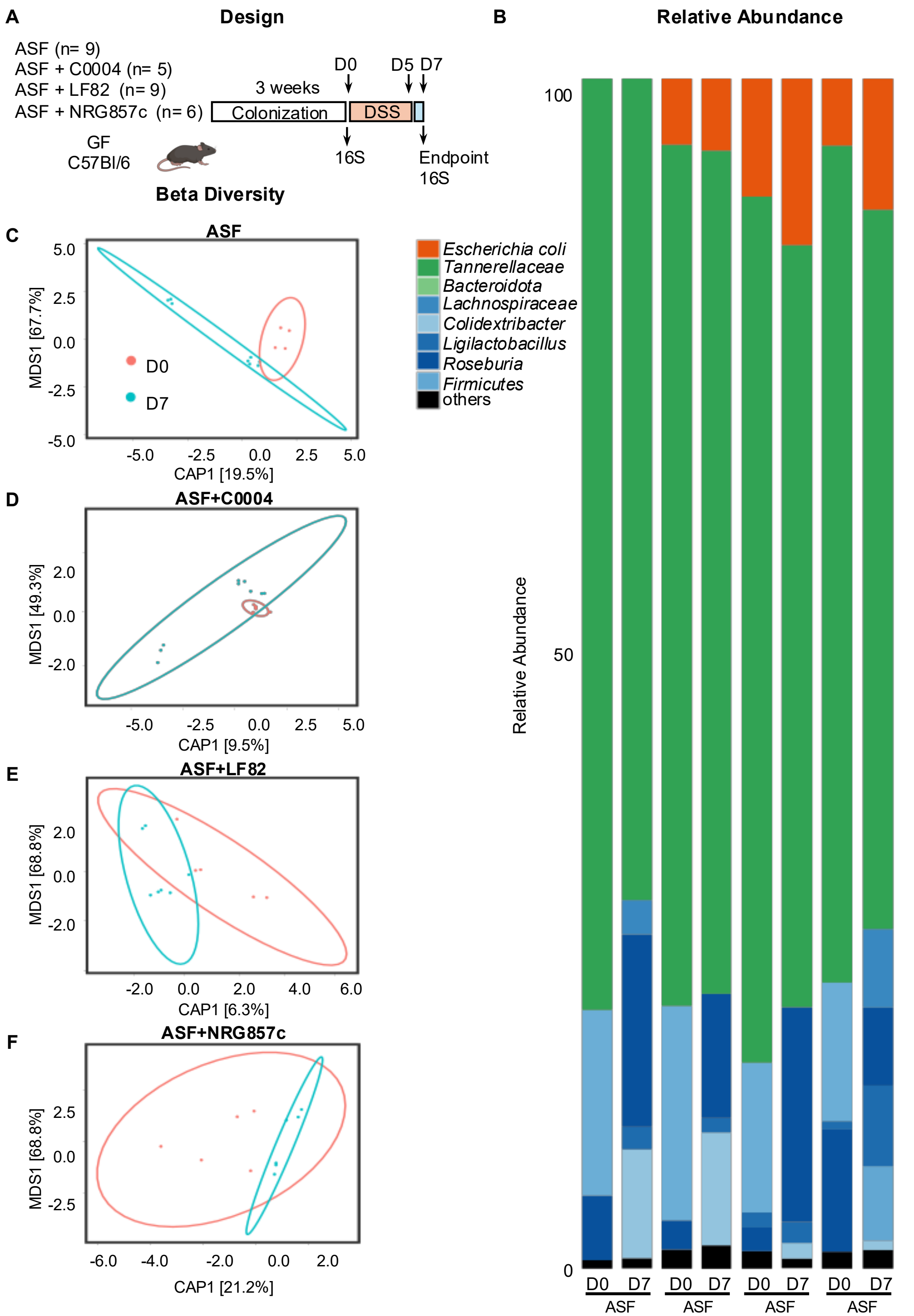

**Fig S1. 16S rRNA microbial profiling before and after DSS colitis.** (**A**) Experimental design. (**B**) Relative abundance plots by group before (D0) and after (D7) DSS induced colitis. Beta diversity plots of (**C**) ASF, (**D**) ASF+C0004, (**E**) ASF+LF82, and (**F**) ASF+NRG857c.

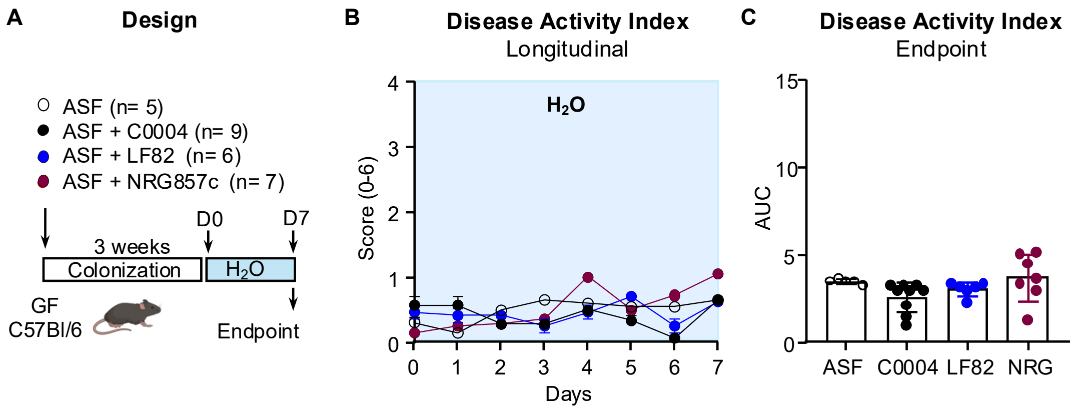


**Fig S2. Gnotobiotic mice in the absence of induced colitis.** (A) Germ-free C57Bl/6 mice were co-colonized with ASF-like and $10^8$ CFU of *E. coli* NRG857c, for 3 weeks. Mice were treated with water. (B) Longitudinal disease activity index (stool consistency + occult blood). (**C**) endpoint area under the curve analysis of disease activity index.

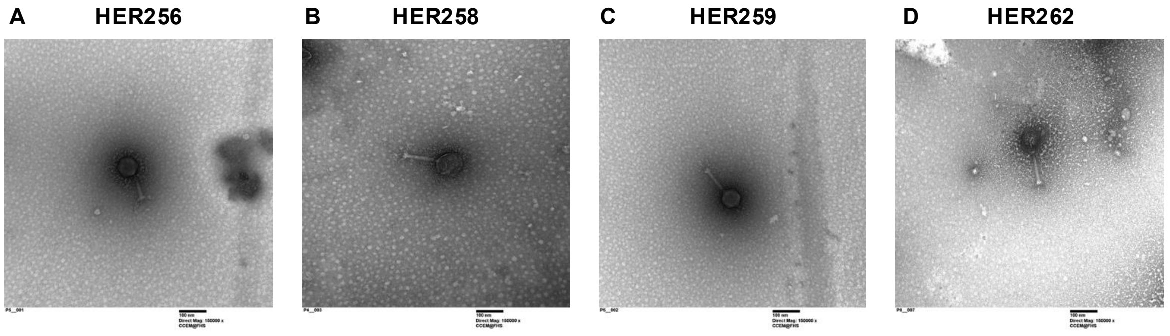


**Fig S3. Transmission electron microscopy of phage evaluated in this study.** (**A**) HER258, (**B**) HER258, (**C**) HER259, and (**D**) HER262

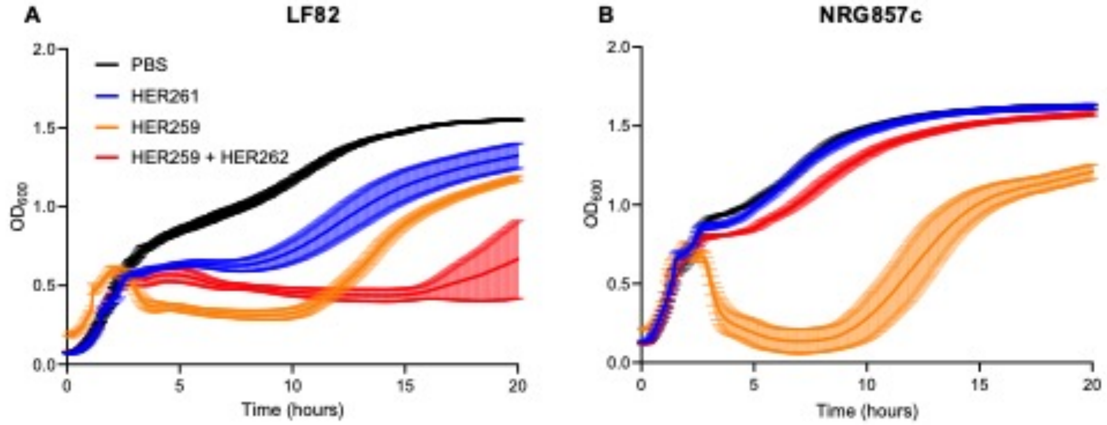


**Fig S4.** HER259 + HER262 cocktail challenges against (**A**) LF82 and (**B**) NRG857c

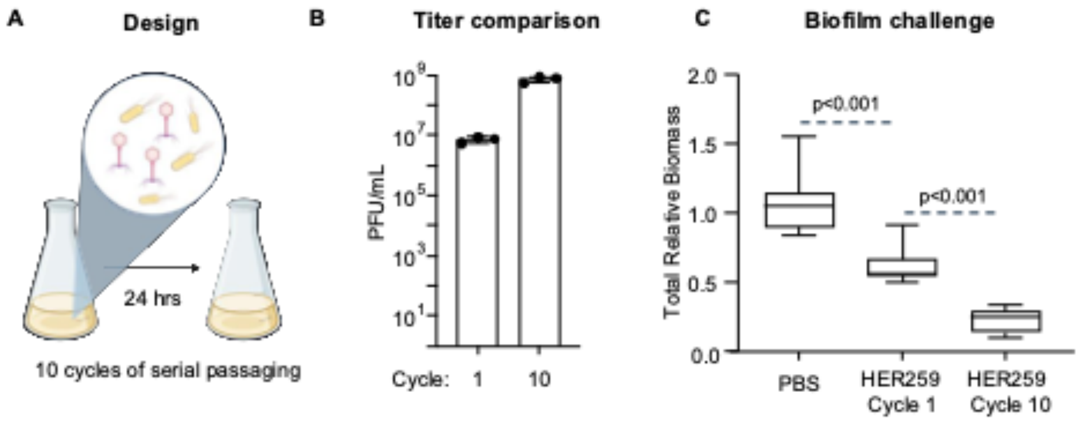


**Fig S5.** (**A**) Serial passaging of HER259 against NR857c. (**B**) HER259 titer between cycle 1 and cycle 10. (**C**) Biofilm HER259 challenge between cycle 1 and cycle 10.

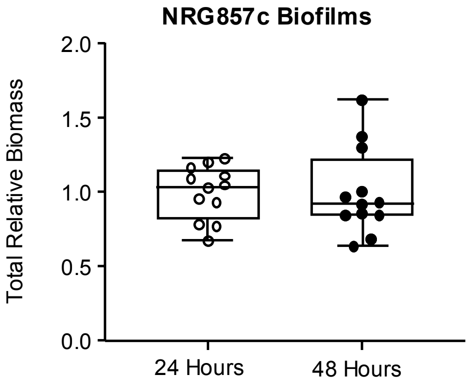


**Fig S6.** Normalized 24- vs 48-hour NRG857c biofilm growth.

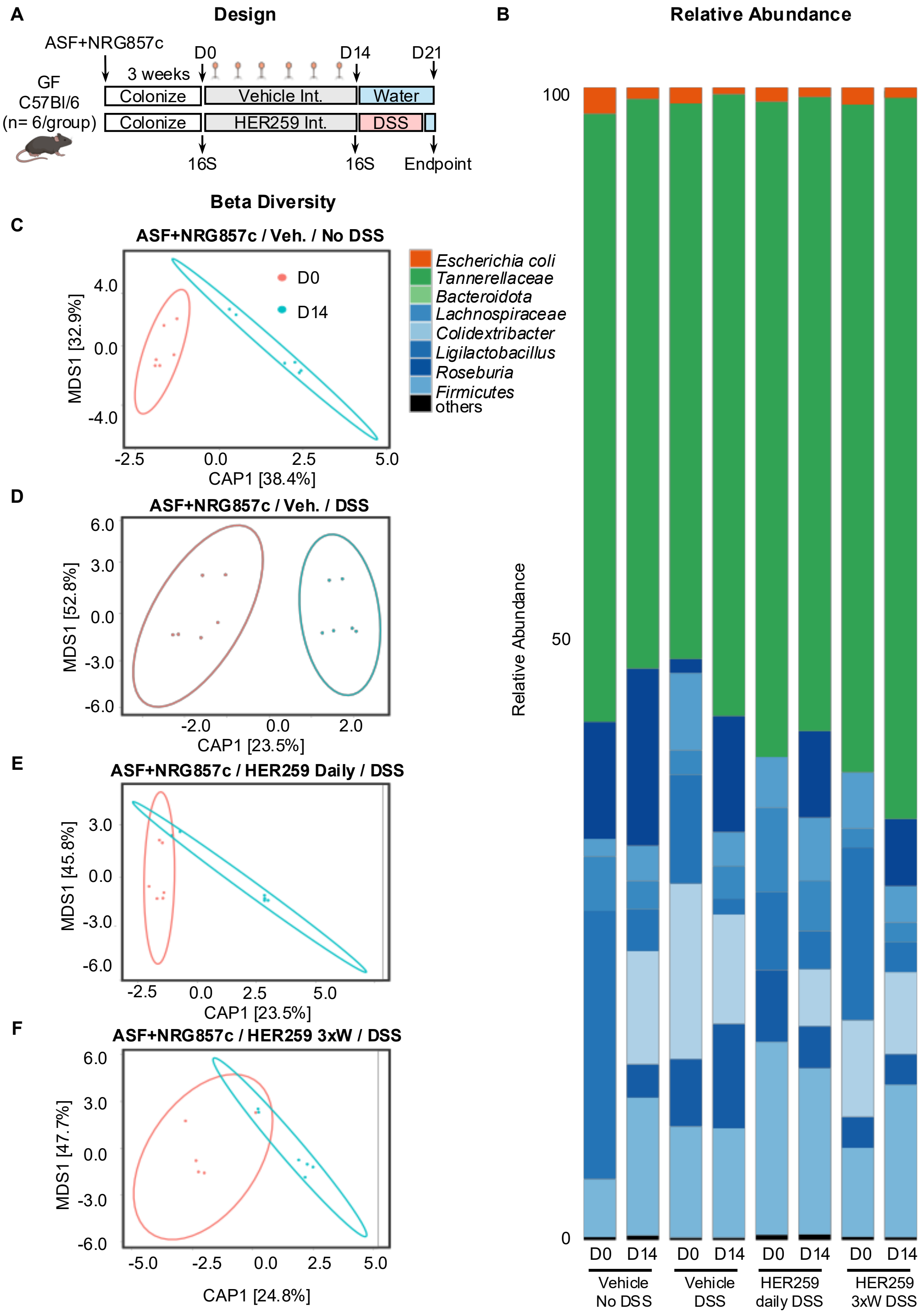
A
Design
ASF+NRG857c
GF
C57Bl/6
(n= 6/group)
3 weeks
D0
D14
D21
Colonize
Vehicle Int.
Water
Colonize
HER259 Int.
DSS
16S
16S
Endpoint
B
Relative Abundance
100
50
0
Relative Abundance
D0
D14
D0
D14
D0
D14
D0
D14
Vehicle No DSS
Vehicle DSS
HER259 daily DSS
HER259 3xW DSS
Escherichia coli
Tannerellaceae
Bacteroidota
Lachnospiraceae
Colidextribacter
Ligilactobacillus
Roseburia
Firmicutes
others
Beta Diversity
C
ASF+NRG857c / Veh. / No DSS
D0
D14
4.0
0.0
-4.0
MDS1 [32.9%]
-2.5
0.0
2.5
5.0
CAP1 [38.4%]
D
ASF+NRG857c / Veh. / DSS
6.0
3.0
0.0
-3.0
-6.0
MDS1 [52.8%]
-2.0
0.0
2.0
CAP1 [23.5%]
E
ASF+NRG857c / HER259 Daily / DSS
3.0
0.0
-3.0
-6.0
MDS1 [45.8%]
-2.5
0.0
2.5
5.0
CAP1 [23.5%]
F
ASF+NRG857c / HER259 3xW / DSS
6.0
3.0
0.0
-3.0
-6.0
MDS1 [47.7%]
-5.0
-2.5
0.0
2.5
5.0
CAP1 [24.8%]

**Fig S7. 16S rRNA microbial profiling before and after phage intervention prior to DSS induced colitis.** (**A**) Experimental design. (**B**) Relative abundance plots by group before (D0) and after (D14) phage intervention. Beta diversity plots of (**C**) ASF+NRG857c mice treated with vehicle, no DSS, (**D**) ASF+NRG857c mice treated with vehicle, DSS (**E**) ASF+NRG857c mice treated with HER259 daily, DSS, and (**F**) ASF+NRG857c mice treated with HER259 3xW, DSS.

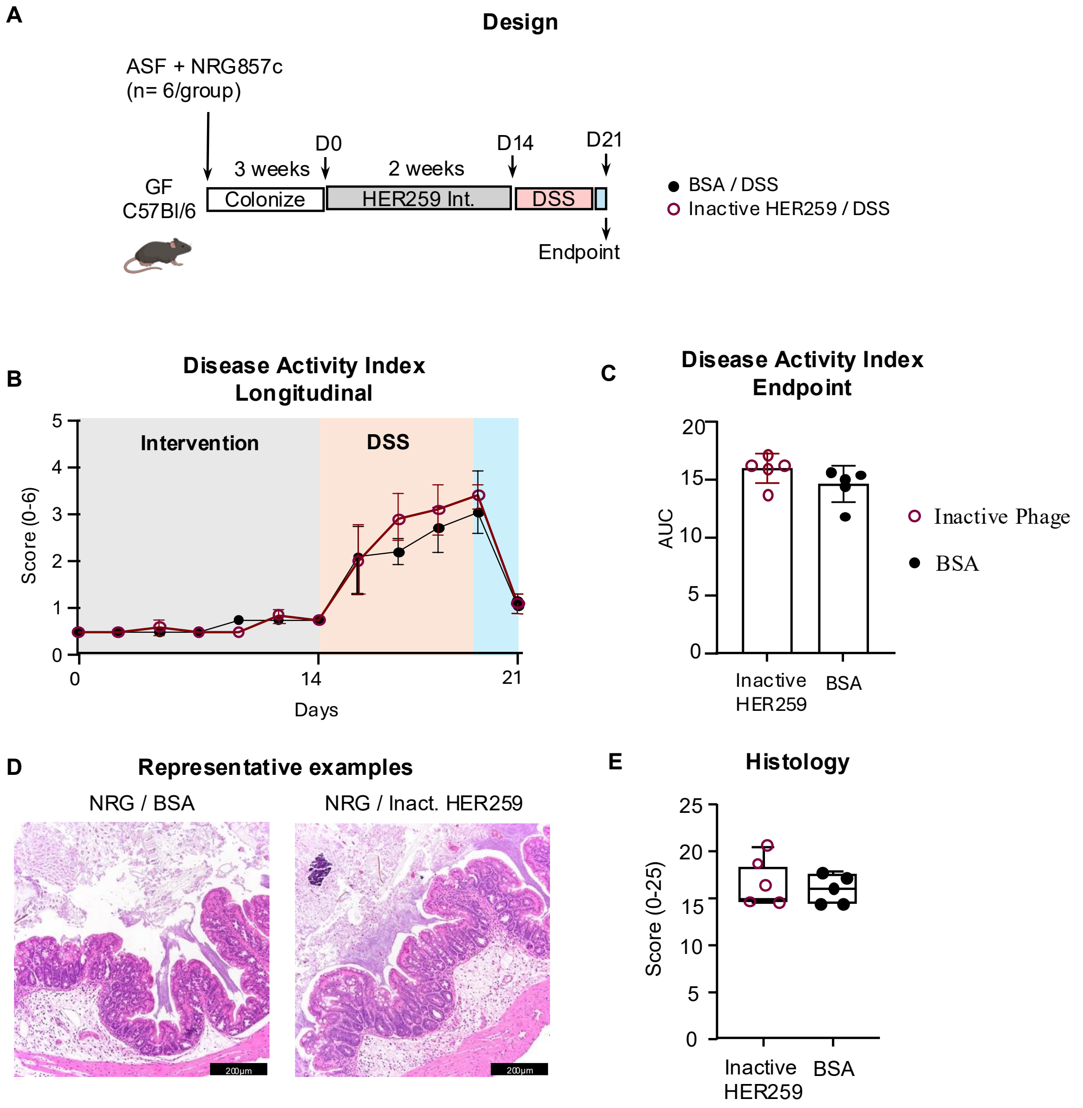


**Fig S8. Heat inactive HER259 does not attenuate bacterial-driven colitis.** (A) Germ-free C57Bl/6 mice were co-colonized with ASF and 108 CFU of E. coli NRG857c, for 3 weeks. Mice were treated for 2 weeks with: 1) BSA (1 mg/mL, PBS with 0.1% bicarbonate); 2) heat inactivated HER259 (1x109 PFU/dose; 0.1% bicarbonate) 3 times/ week. (**B**) Longitudinal disease activity index. (C) endpoint area under the curve analysis of disease activity index. (**D**) Representative endpoint histology of cross-sectional proximal colon taken at 20X. (**E**) Colonic histological scores, determined using a modified pathology score (0-25). Statistical significance determined by Kruskal-Wallis test with Dunn’s post-hoc test or ANOVA with Tukey post-hoc test.

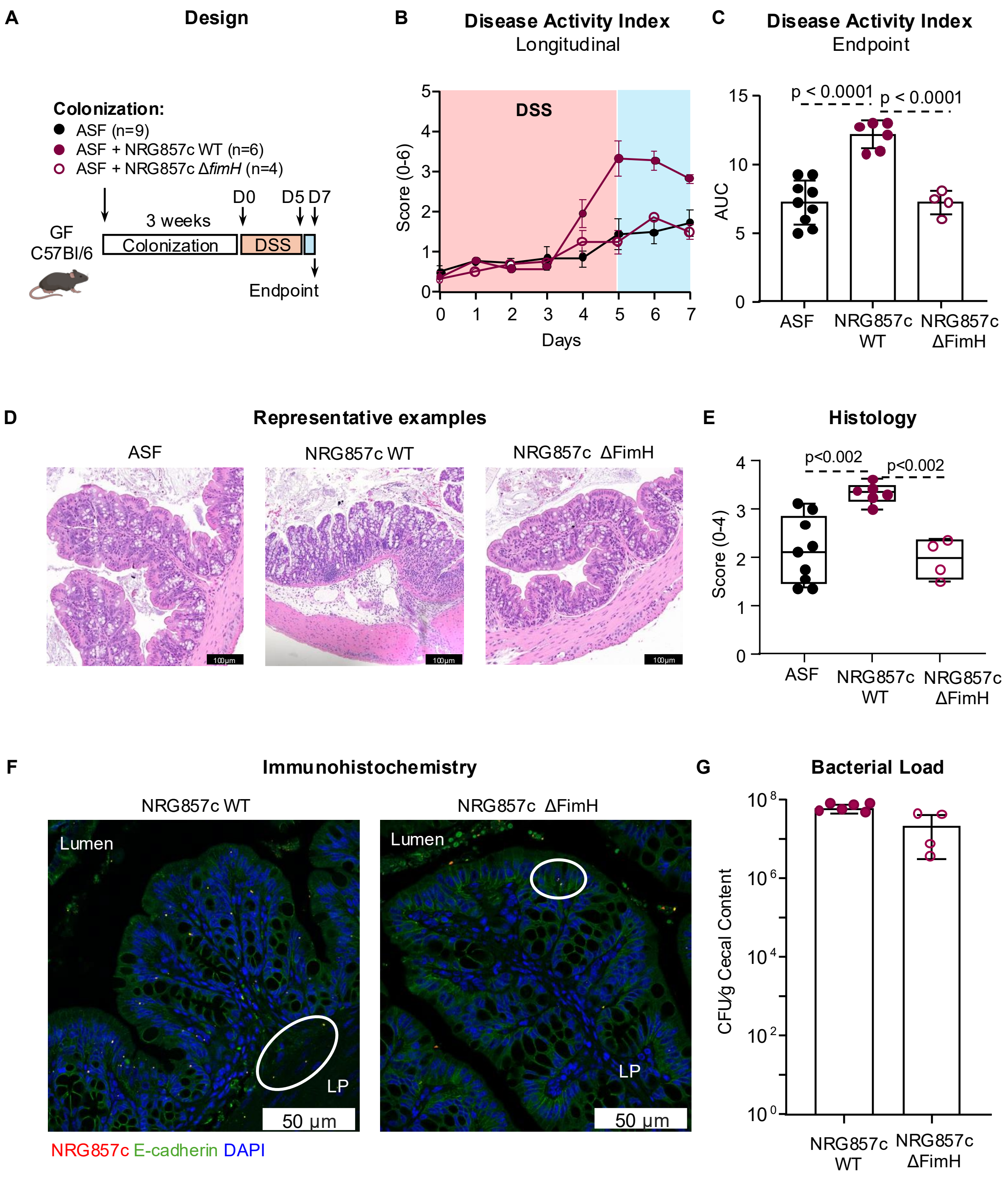


**Fig S9. Gnotobiotic mice colonized with a mutant NRG857c lacking FimH is protected from DSS-induced colitis.** (A) Germ-free (GF) C57Bl/6 mice were colonized with ASF, ASF plus 108 CFU of E. coli NRG857c wild-type (WT) or ASF plus 108 CFU of E. coli NRG857c

ΔFimH for 3 weeks. Mice then received DSS (2%) in drinking water for 5 days followed by 2 days of water. (**B**) Longitudinal disease activity index. (**C**) endpoint area under the curve analysis of disease activity index. (**D**) Representative endpoint histology of cross-sectional proximal colon taken at 40X. (**E**) Colonic histological scores, determined using a modified pathology score (0-25). (**F**) Immunostaining in colonic sections of ASF+NRG857c WT or ASF+NRG857c ΔFimH colonized mice. Yellow represents NRG857c, blue represents DAPI-stained nucleus, and green represents E-cadherin. (**G**) Endpoint cecal bacterial load. Statistical significance determined by Kruskal-Wallis test with Dunn's post-hoc test or ANOVA with Tukey post-hoc test.

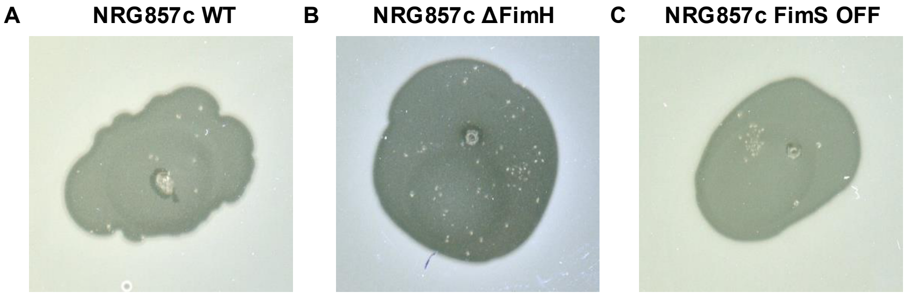


**Fig S10.** HER259 spot test against (**A**) NRG857c WT (B) NRG857c ΔFimH and (**C**) NRG857c FimS OFF.

**Table S1:** Demographics of IMAGINE patients used in this study.

| Characteristics | CD (n=18) | HC (n=11) |
|---|---|---|
| Disease state | | |
| Active | 8 | N/A |
| Remission | 10 | N/A |
| Sex | | |
| Male | 7 | 6 |
| Female | 10 | 5 |
| No info | 1 | 0 |
| Race/ethnicity | | |
| Caucasian | 15 | 9 |
| South Asian | 2 | 1 |
| Other | 1 | 1 |
| Age | 46.07 +/- 17.37 | 56.74 +/- 19.05 |

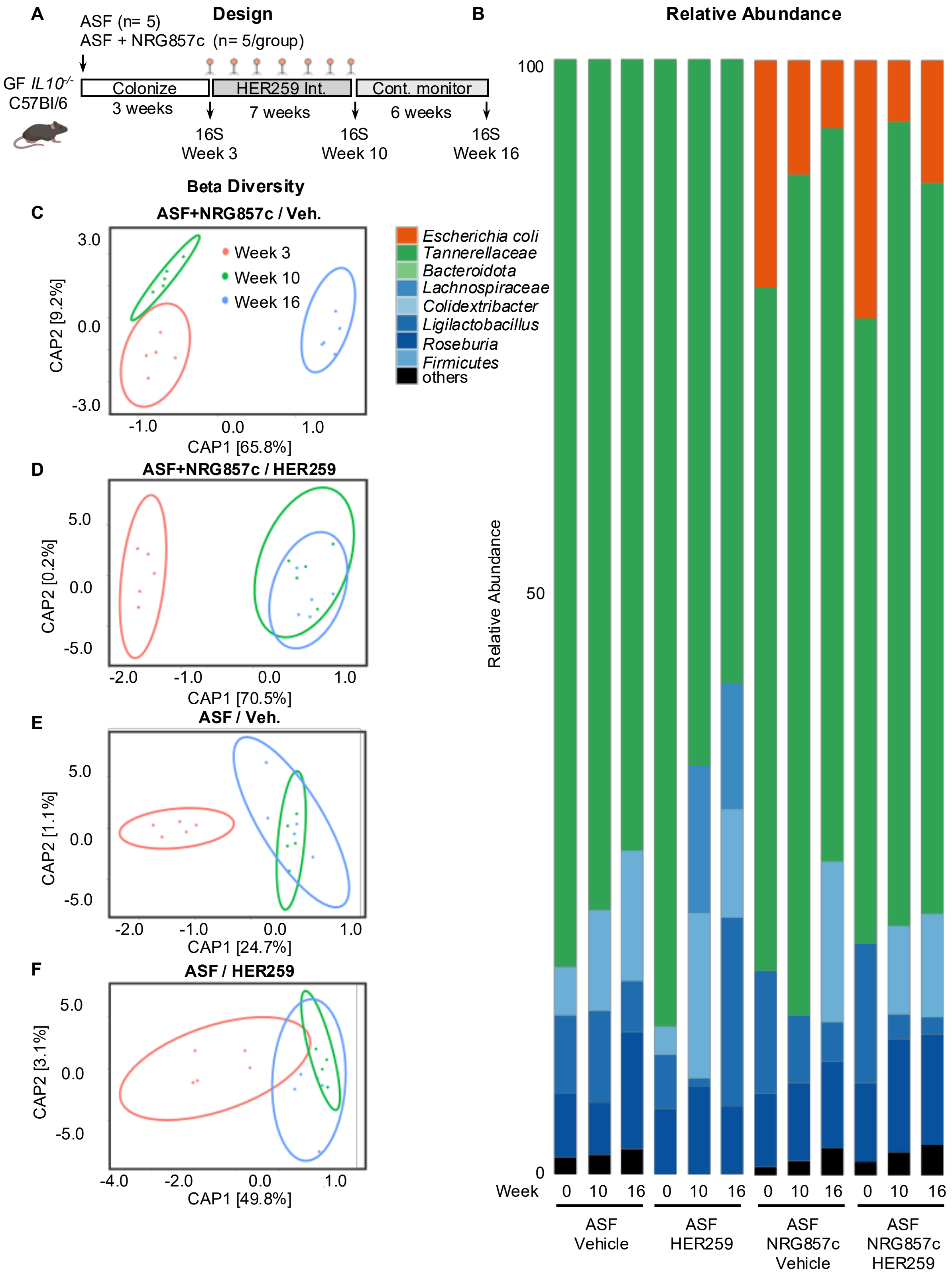
A
Design
ASF (n= 5)
ASF + NRG857c (n= 5/group)
GF IL10-/- C57Bl/6
Colonize
3 weeks
HER259 Int.
7 weeks
Cont. monitor
6 weeks
16S Week 3
16S Week 10
16S Week 16
B
Relative Abundance
Beta Diversity
C
ASF+NRG857c / Veh.
Week 3
Week 10
Week 16
CAP2 [9.2%]
CAP1 [65.8%]
D
ASF+NRG857c / HER259
CAP2 [0.2%]
CAP1 [70.5%]
E
ASF / Veh.
CAP2 [1.1%]
CAP1 [24.7%]
F
ASF / HER259
CAP2 [3.1%]
CAP1 [49.8%]
Escherichia coli
Tannerellaceae
Bacteroidota
Lachnospiraceae
Colidextribacter
Ligilactobacillus
Roseburia
Firmicutes
others
Relative Abundance
Week
ASF Vehicle
ASF HER259
ASF NRG857c Vehicle
ASF NRG857c HER259

**Fig S11. 16S rRNA microbial profiling before and after phage intervention, and six weeks following cessation of phage intervention in IL-10-/- mice.** (**A**) Experimental design. (**B**) Relative abundance plots by group before (Week 3), and after (Week 10) phage intervention, and six weeks following cessation of phage intervention (Week 16). Beta diversity plots of (**C**) ASF+NRG857c mice treated with vehicle (**D**) ASF+NRG857c mice treated with HER259 (**E**) ASF mice treated with vehicle, and (**F**) ASF mice treated with HER259.

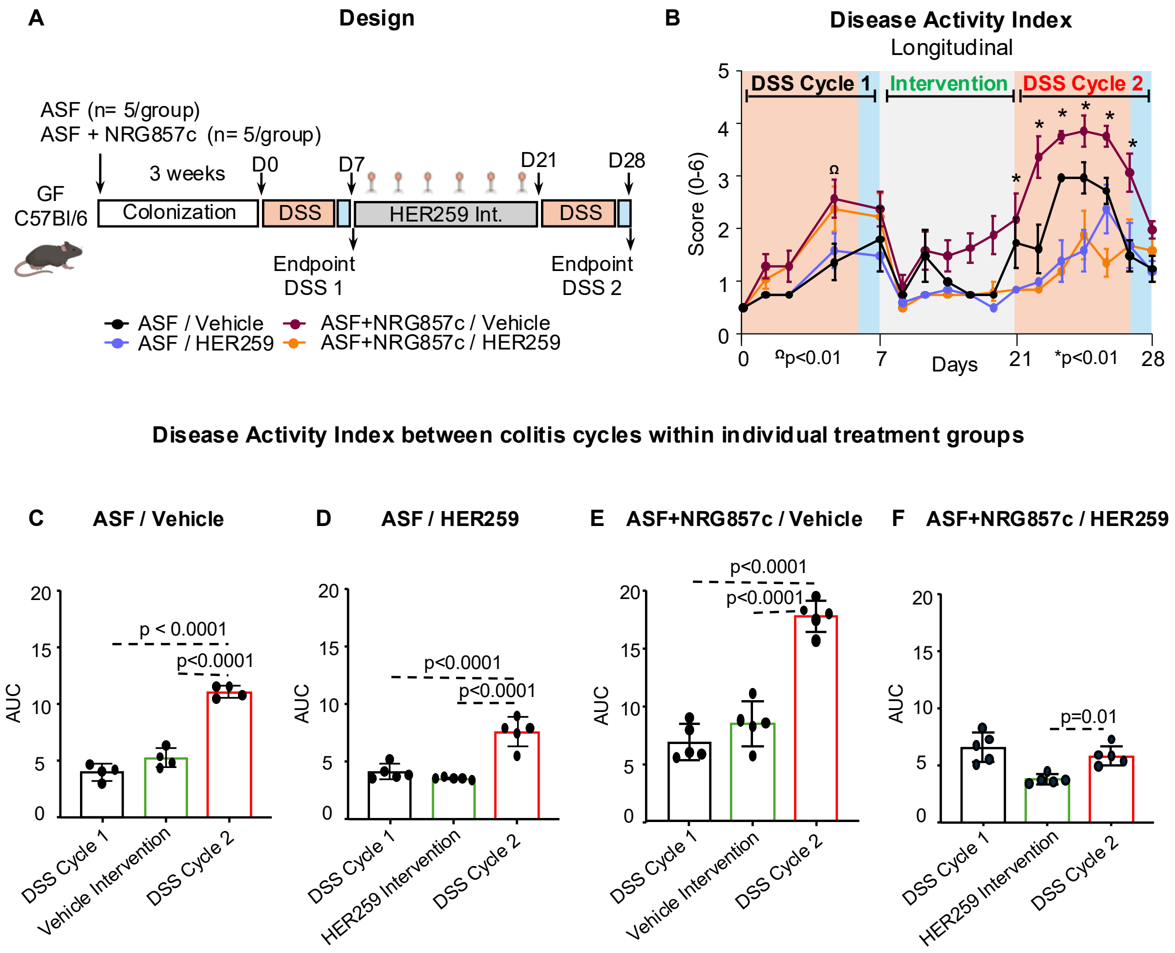


**Fig S12. HER259 treatment prevents reactivation of colitis.** (**A**) experimental design. (**B**) Longitudinal disease activity index (Ωp<0.01 ASF vs. ASF+NRG857c; *p<0.01 ASF+NRG857c/Vehicle vs. ASF+NRG857c/HER259). Segmented AUC analysis of DSS Cyle 1 (black), intervention (green), and DSS Cycle 2 (red) of (**C**) ASF/Vehicle, (**D**) ASF+NRG857c/Vehicle, (**E**) ASF/HER259, and (**F**) ASF+NRG857c/HER259. Statistical significance determined by Kruskal-Wallis test with Dunn's post-hoc test or ANOVA with Tukey post-hoc test.

**Table S2:** qPCR primers used in this study.

| Primer | Sequence (5' → 3') | Orientation | Target | PMID |
|---|---|---|---|---|
| CMD1246 | ACC GTA ACG CAG ACT CAT CCT CAT | Forward | FimS ON | 22665376 |
| CMD1247 | TGA ACG GTC CCA CCA TTA ACC G | Reverse | FimS OFF | |
| CMD1248 | TCA CAT CAC CTC CGC TAT ATG T | Forward or Reverse | | |
| *E. Coli* 16SF | GTT AAT ACC TTT GCT CAT TGA | Forward | E coli 16S rRNA gene | 21440012 |
| *E. Coli* 16SR | ACC AGG GTA TCT AAT CCT GTT | Reverse | | |
| UnivF | TCC TAC GGG AGG CAG CAG TG | Forward | Bacterial 16S rRNA gene | 37131291 |
| UnivR | TTA CCG CGG CTG CTG GCA CG | Reverse | | |
